# Iron-sensing and redox properties of the hemerythrin-like domains of Arabidopsis BRUTUS and BRUTUS-LIKE2 proteins

**DOI:** 10.1101/2024.10.23.619806

**Authors:** Jacob Pullin, Jorge Rodríguez-Celma, Marina Franceschetti, Julia E A Mundy, Dimitri A Svistunenko, Justin M Bradley, Nick E Le Brun, Janneke Balk

## Abstract

Iron uptake in plants is negatively regulated by highly conserved hemerythrin (Hr) E3 ubiquitin ligases exemplified by *Arabidopsis thaliana* BRUTUS (BTS). Physiological studies suggest these are the elusive plant iron sensors, but biochemical evidence is lacking. Here we demonstrate that the N-terminal domains of BTS and BTS-LIKE2 (BTSL2) respectively bind three and two diiron centres within three closely packed Hr-like subdomains. The centres can be reversibly oxidized by O_2_ and H_2_O_2_, resulting in a di-Fe^3+^ form that is non-labile. In the reduced state, a proportion of the iron becomes labile, based on accessibility to Fe^2+^ chelators and reconstitution experiments, consistent with dynamic iron binding. Impaired iron binding and altered redox properties in the BTS *dgl* variant correlate with diminished capacity to suppress the downstream signalling cascade. These data provide the biochemical foundation for a mechanistic model of how BTS/Ls function as iron sensors that are unique to the plant kingdom.

## INTRODUCTION

The concentration of iron in cells and whole organisms is tightly controlled to meet demand while avoiding toxicity from redox cycling of free iron (Fenton chemistry). In plants, the uptake of iron is negatively regulated by specific E3 ubiquitin-protein ligases, known as Hemerythrin RING and Zinc-finger (HRZ) proteins in monocots, and named BRUTUS (BTS) in dicot species^1^. The related BTS-LIKE (BTSL) proteins are found in dicots only and have a specific role in regulating iron uptake in roots^2,3^. Mutations in the rice *HRZ* or Arabidopsis *BTS* genes result in accumulation of excessive amounts of iron, leading to embryo lethality in knockout alleles^4–6^. By contrast, Arabidopsis plants lacking expression of both the *BTSL1* and *BTSL2* genes are viable and accumulate moderate amounts of iron under certain growth conditions^2,3^.

The HRZ/BTS proteins are relatively large (∼140 kDa), consisting of consecutive hemerythrin-like sequences at the N-terminus and a RING-type ubiquitin ligase domain at the C-terminus (Fig. 1a). Despite their name, hemerythrins do not bind heme but a pair of cations, usually iron, and are subdivided into canonical O_2_-carrying hemerythrins and hemerythrin (Hr)-like proteins based on the pattern of the metal-binding His and Glu residues^7^. The E3 ligase domain of HRZ/BTS interacts with transcription factors that orchestrate the iron-deficiency response and targets them for degradation^8,9^ (and reviewed in ^1^). Combination of a Hr-like sequence and E3 ligase is also found in the mammalian F-box/LRR-repeat protein 5 (FBXL5)^10,11^, which has a single Hr-like sequence at the N-terminus and a subunit of a Cullin-type E3 ligase at the C-terminus. In the presence of iron and O_2_, FBXL5 promotes interaction with Iron Regulatory Protein 2 (IRP2), followed by ubiquitination and degradation of IRP2^12^. By analogy, the Hr-like domain of HRZ/BTS has been proposed to sense iron and/or O_2_, but no biochemical studies have been reported to date. Moreover, why there are three Hr-like sequences and how redox activity could influence ubiquitin ligase activity are open questions.

**Fig. 1.**
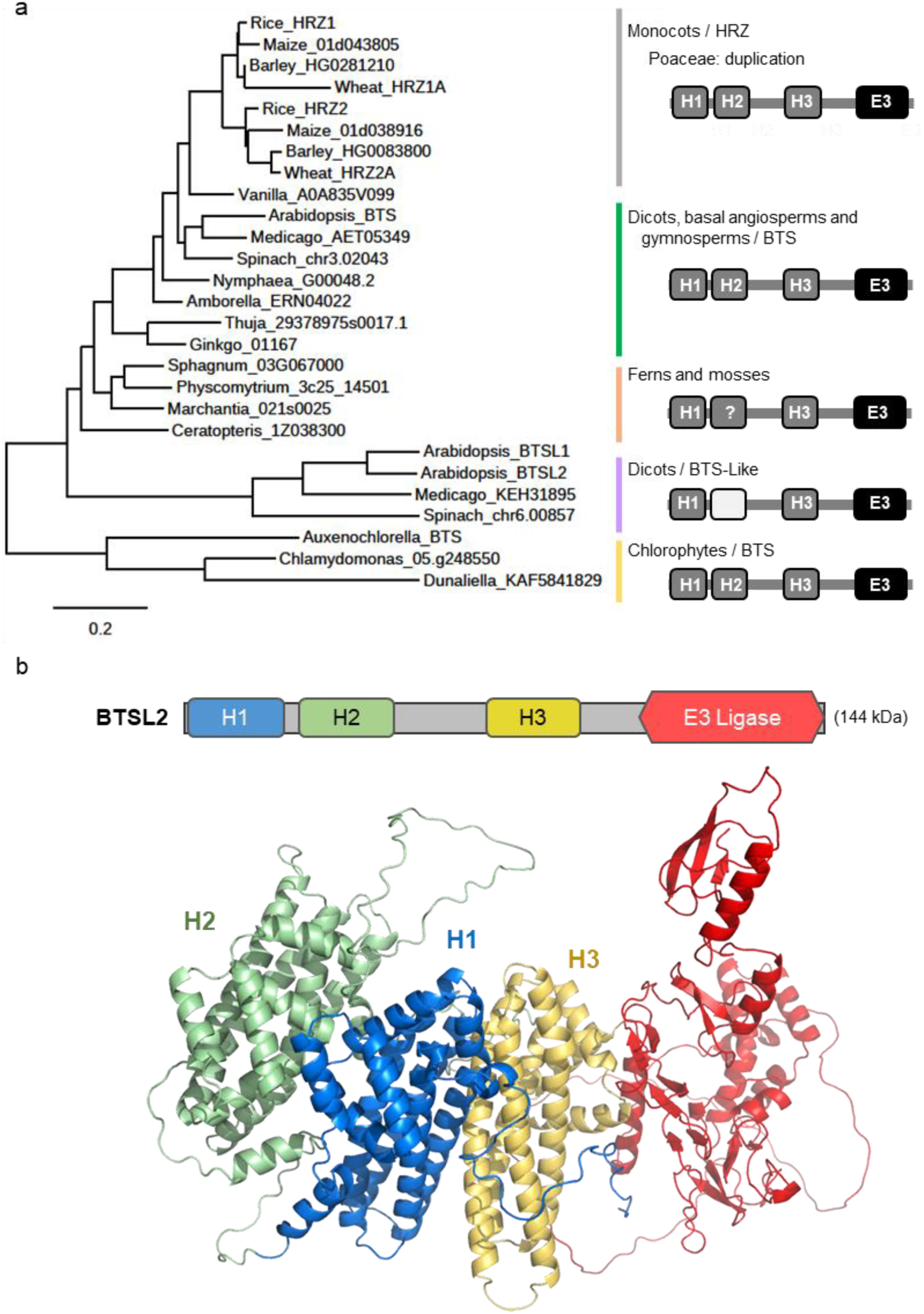
Conserved domain organisation of HRZ/BTS proteins and predicted protein fold. **a** Phylogeny of HRZ, BTS and BTS-Like (BTSL) proteins from selected organisms across the green lineage (left) and their domain organisation (right). The hemerythrin (Hr)-like sequences are marked H1, H2 and H3. Dark grey indicates the presence, and light grey the absence, of iron-binding ligands. The question mark indicates that one of the six ligands is absent or not aligned. For full species names and sequence IDs, see Supplementary Table 1. Phylogenetic analysis was performed using www.phylogeny.fr^16^. The scale bar indicates the number of amino acid substitutions per site. **b** AlphaFold2 model of BTSL2. The Hr-like folds are predicted to be ordered H2–H1–H3 in a tightly packed configuration.

Here we report on the architecture, iron-binding properties and redox activity of the recombinantly expressed Hr-like domains of BTS and BTSL2 from Arabidopsis. Unexpectedly, SAXS analysis showed that the three Hr-like folds are tightly packed in a large, rigid domain, an arrangement supported by AlphaFold2 prediction. A range of spectroscopic methods showed that BTS and BTSL2 bind three and two diiron centres, respectively, which were strongly bound in the oxidized state, but partially labile upon reduction. Impaired iron binding and oxidation activity in vitro were correlated with decreased activity of BTS in vivo. These data suggest a model for the iron-sensing mechanism of BTS/L proteins by means of diiron centres in a unique triple Hr-like domain.

## RESULTS

### The extended N-terminus with three Hr-like sequences is conserved in BTS/L proteins throughout the green lineage

Previous phylogenetic analyses identified HRZ/BTS homologues in different clades of the green lineage, including green algae^1,4,13–15^. Similar to higher plants, the expression of algal homologues is induced in response to iron deficiency in *Chlamydomonas reinhardtii*^13^, *Dunaliella* species and diatoms^15^, suggesting that the role of HRZ/BTS in iron homeostasis is highly conserved. While most HRZ/BTS protein sequences are predicted to have three sequential Hr-like motifs, homologs with only one or two motifs were reportedly found in some plant genomes using bioinformatics^14^. With the ever-expanding availability and completeness of genome sequences, we revisited the phylogenetic analysis to investigate possible variation in the number of Hr-like motifs. All major groups of angiosperms, ferns, mosses and algae have at least one protein of 1200 - 1400 amino acid residues with 50 - 70% overall sequence similarity to Arabidopsis BTS. The phylogenetic relationship of these proteins follows the accepted evolution of the green lineage (Fig. 1a). All the investigated HRZ/BTS sequences had an N-terminal sequence over 950 residues in length encompassing three distinct sequences with a predicted alpha-helical bundle fold typical of Hr-like proteins. This included sequences of HRZ2 in different rice cultivars, indicating that the previously reported open reading frame with one Hr-like sequence was incomplete^4^.

Gene duplication and functional specialization of *HRZ*/*BTS* genes have occurred at least twice: Poaceae (grasses including cereal crops) harbour two *HRZ* paralogues (Fig. 1a). *HRZ1* and *HRZ2* are very similar in sequence but are functionally non-redundant, as shown by mutant studies in rice^4^. Eudicots including Arabidopsis have an additional monophyletic clade of *BTSL* genes, encoding protein sequences that have diverged substantially from BTS (44% similarity). Arabidopsis *BTSL1* and *BTSL2* have more specialized functions in regulating iron uptake in roots^1,8^, although the precise functional differences with *BTS* remain to be established.

In the main HRZ/BTS clade, all three Hr-like sequences contain the diiron-binding motif H-HxxxE-H-HxxxE (InterPro IPR012312). In the BTSL clade, the second sequence has an incomplete H/E motif but is predicted to have the typical alpha-helical bundle fold, as previously noted^1^. In some moss sequences, one of the six H/E residues is lacking or not well-aligned (indicated by a question mark in Fig. 1a).

Models of the HRZ/BTS proteins generated using AlphaFold2 (see examples at www.uniprot.org) consistently showed that the three Hr-like folds are arranged in the order H2-H1-H3 with large contact surfaces between the folds. There is little interaction between the triple Hr-like domain and the E3 ligase domain, except for the direct connection with H3. The predicted structure was similar for the BTSL2 proteins, as depicted in Fig. 1b. This places the metal centres, if present, relatively close to one another (∼21 Å apart) and arranged linearly.

### The three Hr-like folds of BTSL2 form a single inflexible domain structure

To obtain experimental evidence for the AlphaFold2 model of the triple Hr-like domain, the N-terminal part of BTSL2 (BTSL2-N, amino acids 1 - 831) from *Arabidopsis thaliana* was produced in *Escherichia coli* for structural studies. The Maltose Binding Protein (MBP) coding sequence was cloned upstream to improve solubility and a Strep tag was added at the C-terminus for affinity purification (Fig. 2a). Following purification, the MBP-BTSL2-N fusion protein was cleaved with Factor Xa protease and the 98 kDa BTSL-N protein was re-isolated by size-exclusion chromatography (SEC) (Fig. 2b). Because crystallisation trials were unsuccessful, structural data were obtained through small-angle X-ray scattering (SAXS). This low-resolution technique provides information on the average size and shape of proteins in solution^17^. Scattering is measured as the protein elutes from a SEC column (SEC-SAXS) in a series of frames, to enable the selection of the highest quality data, avoiding misfolded and aggregated protein in the sample.

**Fig. 2.**
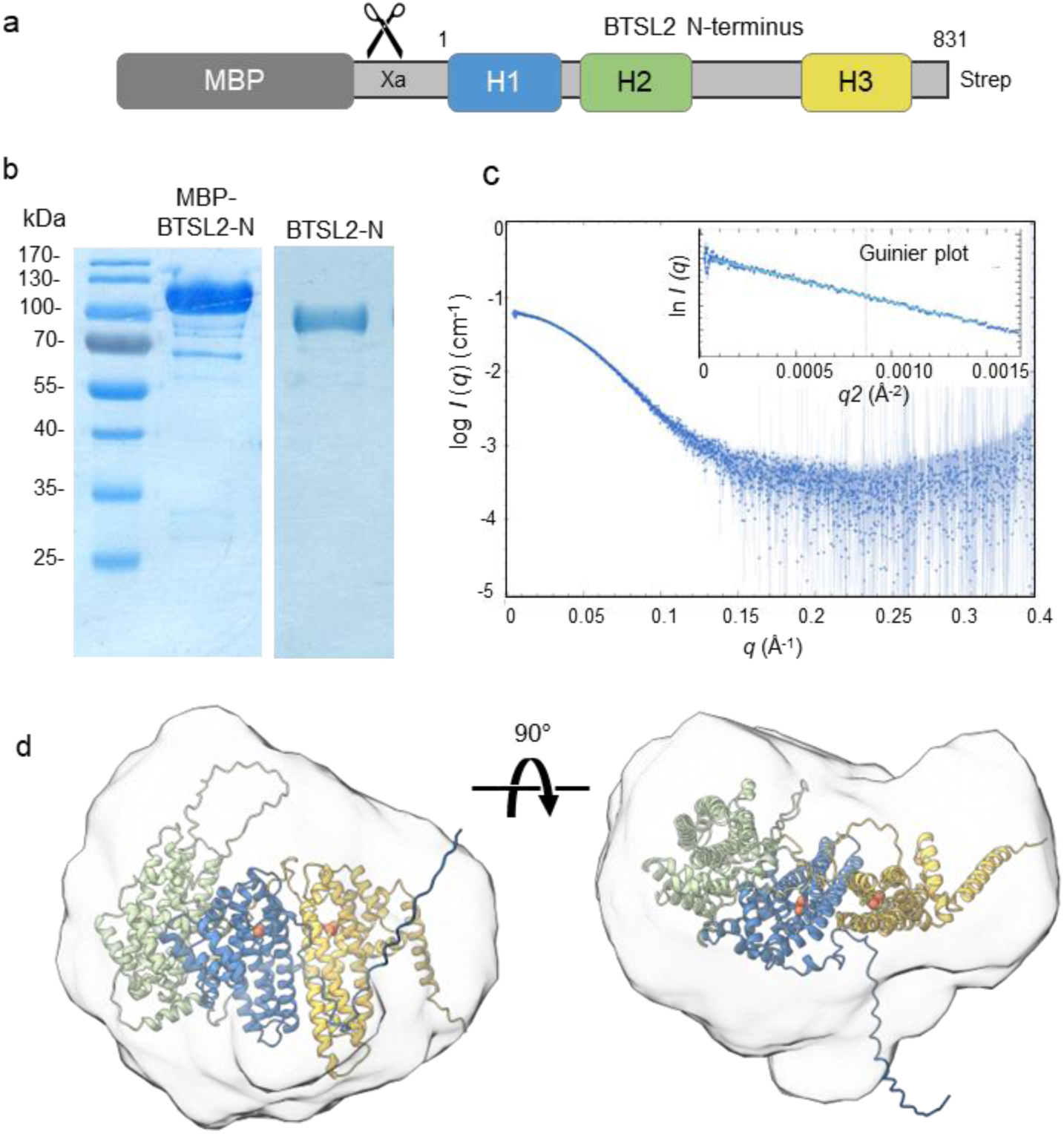
SAXS analysis of the N-terminal domain of BTSL2. **a** Expression construct of the Arabidopsis BTSL2 N-terminus (BTSL2-N, codons 1 – 831) encompassing the three Hr-like sequences, flanked by Maltose Binding Protein (MBP) for solubility and a C-terminal Strep tag for affinity purification. **b** Coomassie-stained SDS-PAGE gel of MBP-BTSL2-N purified using Strep-Tactin XT (left) and of BTSL2-N after removal of the MBP domain following protease Xa cleavage and size-exclusion chromatography (right). The images are representative of three experiments. **c** Scattering profile of intensity *I*(*q*), presented in log-scale, versus *q*, the momentum transfer variable which is inversely correlated with the X-ray wavelength in Ångstrom (Å^-1^). Inset: Guinier analysis, in which ln(*I*) is plotted against *q*^2^. A radius of gyration *Rg* of 33.95 ± 0.05 Å was calculated from the slope of the graph (R^2^ = 0.9941). The intensity at zero angle *I*(0) was 0.062 cm^-1^ ± 0.000096 and the residual values showed an even distribution. **d** Model of the BTSL2-N protein envelope overlaid with a refined AlphaFold2 model with manually inserted diiron centres. See Supplementary Data 1 for a summary of SAXS experimental data and analysis. Source data have been deposited under accession SASDU79 in www.sasbdb.org.

Four frames in the middle of the single elution peak (Supplementary Fig. 1a) were selected and merged to give the scattering profile of intensity (*I*) as a function of *q*, the momentum transfer variable (Fig. 2c). The shape of the profile was typical of non-spherical monodispersed particles^17^. Plotting the natural logarithm of intensity, ln(*I*), versus the square of lower *q* values, known as Guinier analysis, gave a straight line which confirmed the homogeneity of the sample (Fig. 2c, inset). The slope of the Guinier plot was used to calculate a radius of gyration *R*_g_ of 33.95 ± 0.05 Å. The Bayesian inference method^18^ gave a molecular mass of 101 kDa, which is close to the expected molecular mass of 97.771 kDa calculated from the amino acid sequence (Supplementary Data 1) and shows that the recombinant protein is monomeric. The shape of the Kratky plot of *q*^2^ x *I* versus *q* (Supplementary Fig. 1b) indicated that the particle is globular or globoid with only a small degree of flexibility, in agreement with tight packing of the Hr-like folds.

SAXS data can also be assessed in real space using the pair distance distribution function, *P*(*r*), which is obtained by indirect Fourier transformation of *I*(*q*) (Supplementary Fig. 1c). This analysis gave a maximum distance (*d*_max_) of 98 Å, in agreement with the longest predicted axis of 96.5 Å in the AlphaFold2 model. The protein envelope calculated from raw scattering data using DAMMIN software^19^ is shown in Fig. 2d, overlaid with a refined AlphaFold2 model. The *χ*^2^ value of the fit between model and envelope was 1.075, or 1.008 for the model to the scattering data using CRYSOL analysis (Supplementary Data 1). From this we can conclude that the AlphaFold2 structure is representative of the solution structure of BTSL2-N as determined by SAXS.

### The Hr-like domains of BTSL2 and BTS bind diiron centres, except for H2 in BTSL2

To determine the type and number of metal cofactors in the N-terminal part of BTSL2 and BTS, the purified proteins were analysed by Inductively Coupled Plasma – Mass Spectrometry (ICP-MS) followed by quantitative assays to determine the molar ratio of iron per polypeptide. The MBP domain was not removed to maximize protein yield for these and subsequent analyses (Fig. 3a-b). For BTS-N, addition of β-mercaptoethanol or glutathione to all buffers was required to prevent precipitation. ICP-MS for a range of transition metals showed that iron was approximately nine times more abundant than zinc or manganese associated with purified MBP-BTS-N, and three times more abundant in purified MBP-BTSL2-N (Supplementary Table 2). Using sulfur to estimate the protein concentration, the iron-to-protein ratio was 2.2 ± 0.1 for BTS and 3.0 ± 0.05 for BTSL2. These numbers are thought to be an underestimation, particularly for BTS, because the sample likely contained residual glutathione.

**Fig. 3.**
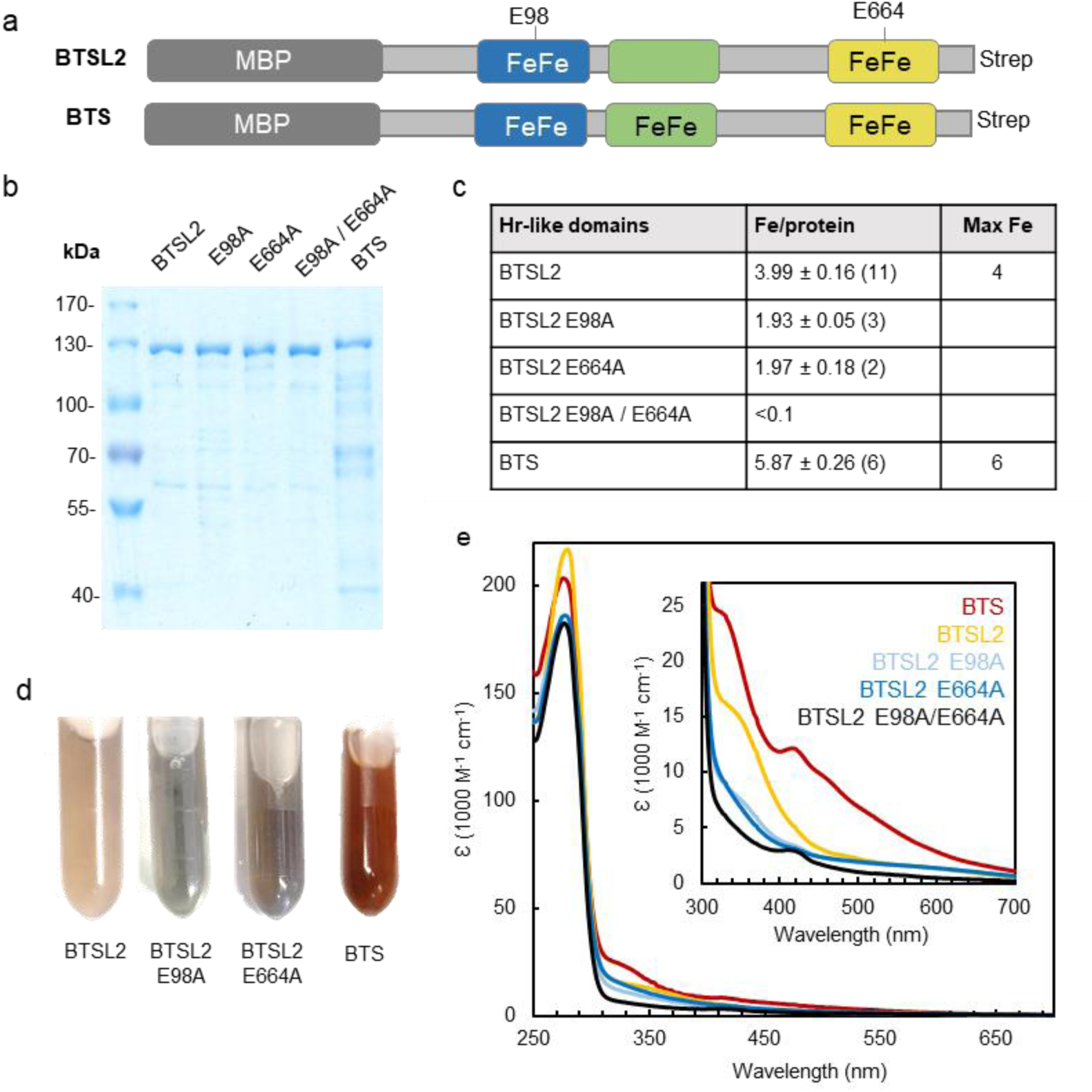
Iron cofactors in the Hr-like domains of BTSL2 and BTS. **a** Diagrams of the MBP fusion proteins of Arabidopsis BTSL2 and BTS N-terminal sequences used for iron quantification and spectroscopy. The E98 and E664 residues are indicated. **b** Coomassie-stained SDS-PAGE gel of the indicated purified proteins including the MBP domain (omitted from the labels), 5 µg protein per lane. The results are representative for the number of preparations given in (c). **c** Quantification of bound iron atoms per polypeptide using the colorimetric chelator Ferene. Values are the mean ± SD of independent protein purifications with n given in brackets. **d-e** Colour (d) and UV-visible spectra (e) of the as-isolated proteins from (a). The protein concentrations were BTSL2, 462 µM; BTSL2 E98A, 345 µM; BTSL2 E664A, 527 µM; BTS, 222 µM in (d) and diluted for UV-visible spectroscopy.

Quantitative analysis of iron using a colorimetric reagent and the Bradford assay for protein showed that MBP-BTSL2-N contained 3.99 ± 0.16 iron atoms per polypeptide (Fig. 3c), close to the expected ratio of 4 based on the two predicted diiron binding sites in H1 and H3 (Fig. 1a). The as-isolated protein was yellow in colour with absorbance around 350 nm (Fig. 3d-e), which is typical of diiron proteins.

To confirm the location of the iron-binding sites in BTSL2-N, we substituted glutamate (E) in the centre of the two H-HxxxE-H-HxxxE motifs with alanine (A) by site-directed mutagenesis. This Glu residue bridges the two iron atoms and E-to-A substitution is known to completely abolish iron binding at hemerythrin sites^11,20^. The E98A substituted protein was isolated with approximately two iron per polypeptide, and a similar result was obtained with E664A (Fig. 3a, c). The single variant proteins were greyish in colour (Fig. 3d) and had lost most of the absorbance in the 350 nm region (approximately 8000 M^-1^ cm^-1^, Fig. 3e). Combining E98A and E664A decreased the amount of iron associated with BTSL2-N below the detection limit of the colorimetric chelator (Fig. 3c) despite some residual absorbance (Fig. 3e).

Purified BTS-N contained 5.87 ± 0.26 iron atoms per polypeptide (Fig. 3c), close to the expected ratio of 6 for three diiron centres, one in each Hr-like subdomain. The as-isolated protein was reddish in colour giving a distinct UV-visible spectrum with features at 332 and 420 nm (Fig. 3d-e). These are in part reminiscent of an iron-sulfur (Fe-S) cluster, but further spectroscopic analysis ruled out this possibility (see below).

These data, together with the structure of the N-terminus, showed that BTSL2 lacks a diiron centre in the distally-located H2 motif and thus has two aligned diiron centres proximal to the E3 ligase domain. BTS has a full complement of three diiron centres.

### The Hr-like iron centres react with O_2_ and H_2_O_2_

Next, we investigated whether the Hr-like domains of BTSL2 and BTS are redox-active. The UV-visible spectra of BTSL2-N in Fig. 3e indicated that the diiron centres are predominantly in the oxidized, diferric state in the ‘as-isolated’ protein. Attempts to obtain proteins with diiron centres in a stable reduced state under atmospheric conditions failed, and therefore samples were transferred to an anaerobic chamber where they were mixed with a small excess of sodium dithionite as reductant, followed by buffer exchange on a desalting column to remove dithionite.

To probe the reaction with O_2_ and H_2_O_2_, reduced BTSL2-N was titrated with incremental aliquots of oxidant and changes in UV-visible absorbance were monitored (Fig. 4a-b). For both O_2_ and H_2_O_2_ this resulted in recovery of the absorbance signals of the as-isolated protein. Changes in the circular dichroism (CD) spectrum showed similar behaviour (Supplementary Fig. 2), indicating that BTSL2-N was completely re-oxidized without loss of iron cofactor. Reduction of BTS-N resulted in a less pronounced decrease in absorbance at 320 – 400 nm than for BTSL2-N. Addition of O_2_ or H_2_O_2_ returned the absorbance signals to those of the ‘as-isolated’ spectra (Fig. 4d-e).

**Fig. 4.**
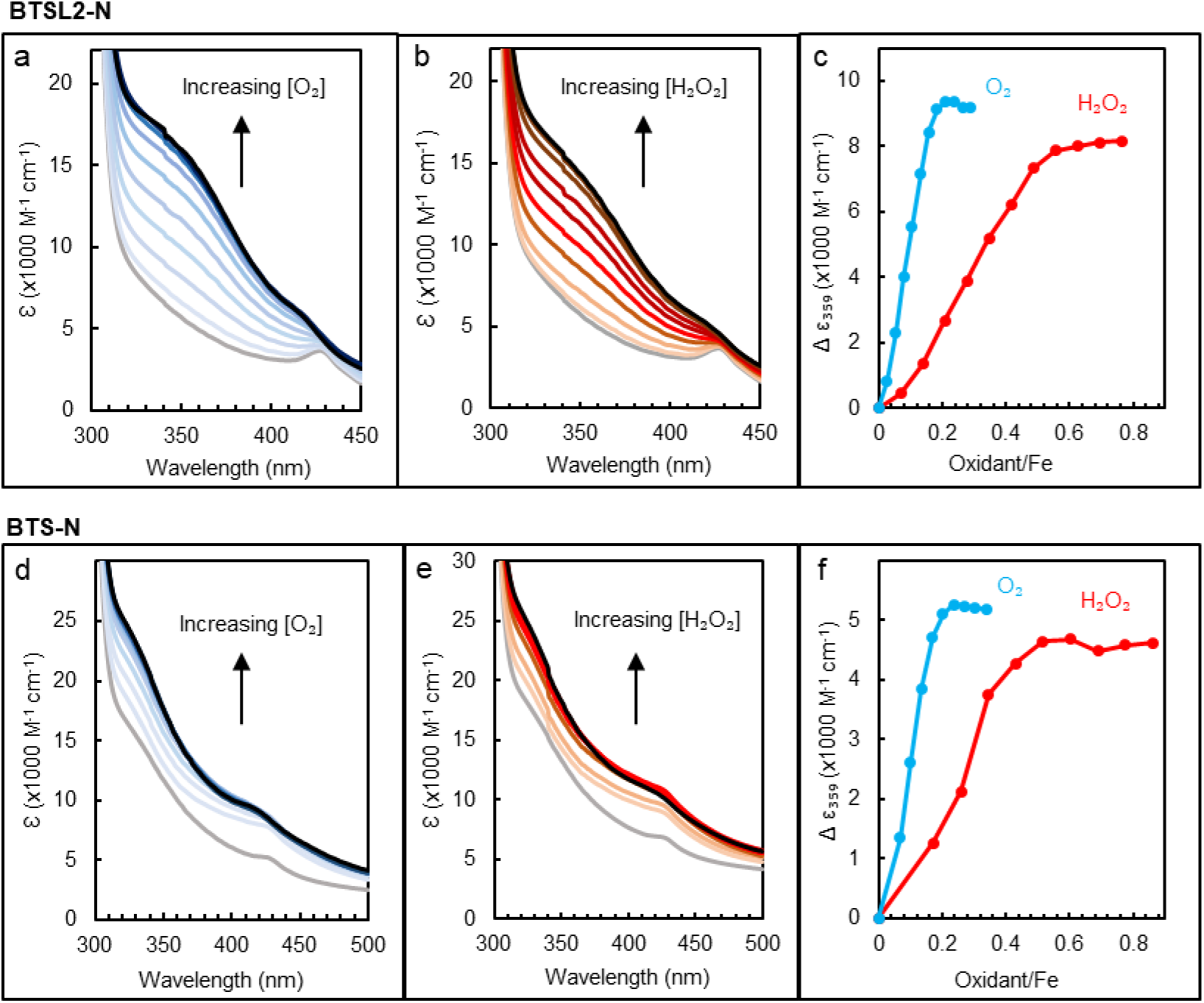
The Hr-like domains of BTSL2 and BTS are redox-active with O2 and H2O2. Redox titrations monitored by UV-visible absorbance spectroscopy of (**a-b**) BTSL2-N and (**d-e**) BTS-N (MBP fusion proteins, see Fig. 3). Black line: ‘as-isolated’ proteins purified in air (a, d) or end point of the titration (b, e). Grey line: fully reduced with dithionite in an anaerobic chamber and desalted. Colour lines: titration with O2 (a and d, blue lines) or H2O2 (b and e, orange-red lines). **c, f** The difference in absorbance at 359 nm as a function of the oxidant to iron ratio based on the spectra in (a-b) for BTSL2-N, and spectra in (d-e) for BTS-N.

The observed stoichiometries of oxidant to iron were ∼0.2 for O_2_ and ∼0.5 for H_2_O_2_ (Fig. 4c, f), close to the theoretical minima of 0.25 for O_2_ and 0.5 for H_2_O_2_. The ratios are typical for reactions of diiron centres, such as Dps mini-ferritins, bacterioferritin, ribonucleotide reductase or alternative oxidase^21–24^, with O_2_ and H_2_O_2_ accepting four and two electrons, respectively. To investigate whether the O_2_ reaction proceeds via a H_2_O_2_ intermediate, we probed for H_2_O_2_ formation using the colorimetric dye Amplex Red. Samples of reduced BTSL2-N and BTS-N protein containing 40 µM Fe^2+^ were mixed with air-saturated Amplex Red assay reagents. Following reaction with 20 µM O_2_, the theoretical maximum production of H_2_O_2_ would be 20 µM, assuming that all H_2_O_2_ formed reacted with the Amplex Red reagent. For BTSL2-N and BTS-N, 24.3 and 15.6 µM H_2_O_2_ were detected, respectively. We also tested if glutathione could reduce the diiron centres in place of dithionite. A physiological concentration of glutathione (3 mM) was mixed with anaerobic BTSL2-N, and we observed reduction of the protein, albeit at a very slow rate (Supplementary Fig. 3).

Because the stoichiometry for the reaction with O_2_ was somewhat lower than expected, we investigated the possibility that the diiron centres in BTSL2-N and BTS-N were not fully reduced in the presence of excess dithionite, resulting in a proportion of semi-reduced, paramagnetic centres. Electron Paramagnetic Resonance (EPR) spectroscopy of the as-isolated proteins showed a prominent signal at g = 4.3, which was essentially absent from the spectra of the reduced proteins (Fig. 5a-b). The signal corresponds to mononuclear Fe^3+^ but represented less than 1% of the total iron in the samples (see quantification described in Materials and Methods). Both BTSL2-N and BTS-N proteins also have an anisotropic EPR signal with g-values in the ≈1.9 area, which can be attributed to a mixed-valent Fe^2+^/Fe^3+^ centre with a µ-oxo bridging ligand rather than a µ-hydroxo-bridged species, which typically has higher g-values^25^. Again, quantification indicated that this species represented less than 1% of the total iron. Titrations of oxidised protein with dithionite did not significantly increase the amplitude of the mixed valent signals, with only a moderate increase of the signal in BTSL2-N and no change in BTS-N (Supplementary Fig. 4a-b). Interestingly, the mixed valent signal was broader and diminished in the E98A variant of BTSL2-N, associated with the H3 diiron centre that remained (Supplementary Fig. 4c).

**Fig. 5.**
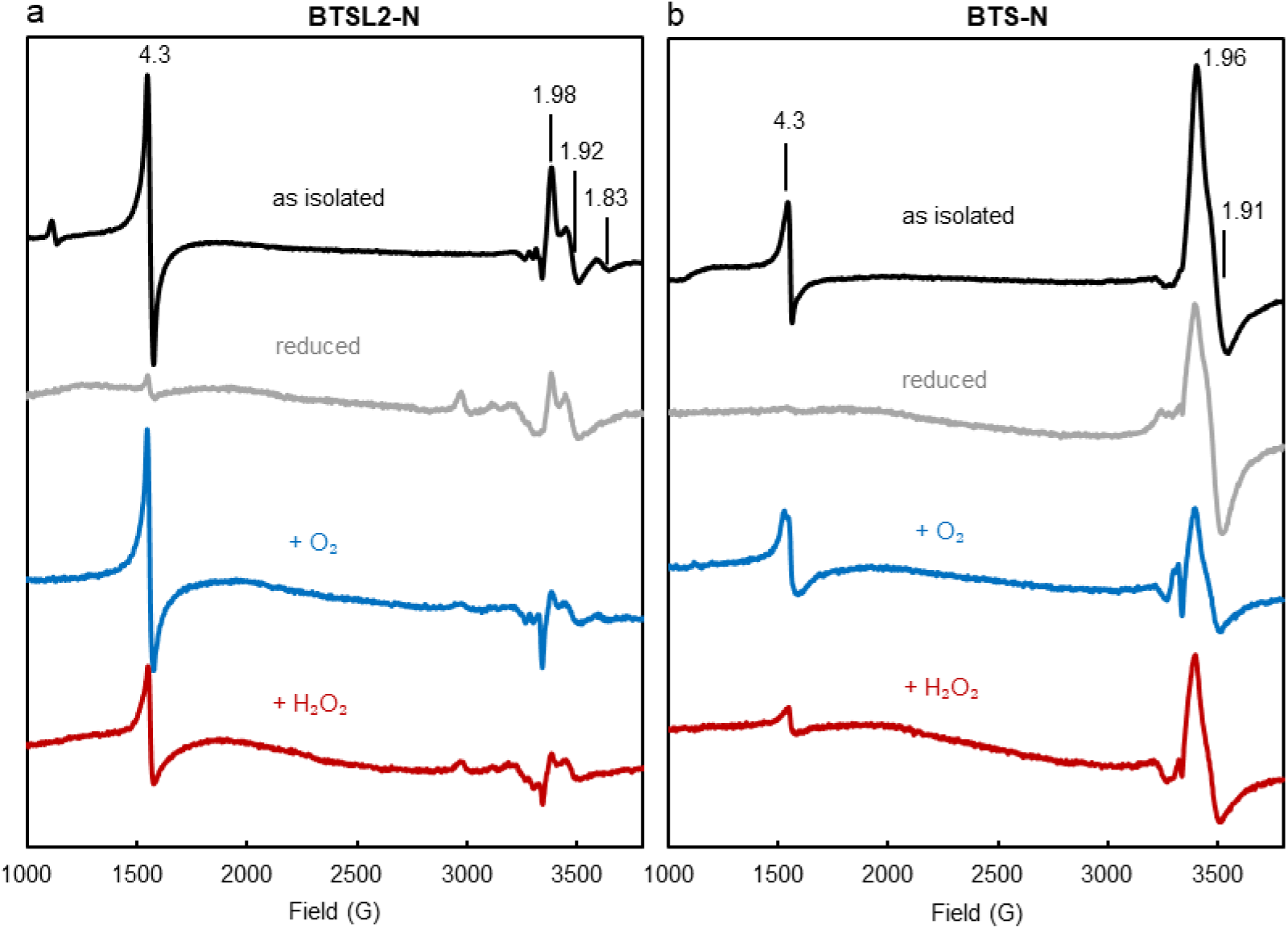
Analysis of protein-bound iron in the Hr-like domains of BTSL2 and BTS by electron paramagnetic resonance (EPR). EPR spectra of (**a**) BTSL2-N and (**b**) BTS-N. The protein domains were expressed as MBP fusion proteins, see Fig. 3a. The signal with g-value 4.3 is mononuclear iron; the signals with g values around 1.9 are characteristic of a mixed-valent Fe^2+^/Fe^3+^ centre. Both signals represent ∼1% of the protein-bound iron, because the majority of the diiron centres are EPR silent when fully oxidized or reduced. Black: BTSL2-N (287 µM) and BTS-N (222 µM) proteins as isolated. Grey: BTSL2-N (130 µM) and BTS-N (100 µM) reduced with dithionite and desalted in an anaerobic chamber. Blue: BTSL2-N (106 µM) and BTS-N (84 µM) 3 min after mixing with calculated volumes of supersaturated O2 solution for end concentrations of 216 µM and 262 µM O2, respectively. Red: BTSL2-N (106 µM) and BTS-N (84 µM) 3 min after mixing with 216 µM and 262 µM H2O2, respectively. Spectra were adjusted to compensate for dilution by reagent additions and desalting. EPR spectra were recorded at 10 K using the following parameters: microwave frequency vMW = 9.353 GHz, modulation frequency vM = 100 kHz, microwave power PMW = 3.17 mW, modulation amplitude AM = 5 G. The results are representative of two independent protein preparations.

The overall lack of signal for the majority of the iron is consistent with the expectation that the fully oxidized diiron centres are EPR silent due to antiferromagnetic coupling of the two S = 5/2 Fe^3+^ ions, which are reduced without accessing the mixed valence Fe^2+^/Fe^3+^ form. For BTS-N, there was no specific signal in the EPR spectrum of the reduced protein with g-values typical of Fe-S clusters^26^. CD spectra of BTS/L proteins were also not consistent with those of known Fe-S proteins^27–29^.

Upon re-exposure to O_2_, the g = 4.3 mononuclear iron signal returned to an intensity similar to that of the as-isolated proteins (Fig. 5). By contrast, exposure to H_2_O_2_ resulted in a signal of much lower intensity. The mixed-valent signal was not responsive to oxidant within the timescale of observation, suggesting that this is not a physiologically relevant species. A small EPR signal at g ≈ 2 had features consistent with protein-based peroxyl radicals and was seen inconsistently among some samples; a spectrum of this signal in BTSL2-N is reported with a narrow field range in Supplementary Fig. 5.

In summary, the data demonstrated that the Hr-like domains of BTSL2 and BTS contain diiron centres that are redox-active with both O_2_ and H_2_O_2_.

### Iron binding to the Hr-like domains of BTSL2 and BTS is reversible in the reduced state

For BTS/L proteins to function as iron sensors, it is expected that they bind iron dynamically. However, iron remained bound to the Hr-like domains of BTSL2 and BTS under the conditions used for purification (Fig. 3) and following reduction and re-oxidation (Fig. 4). Therefore, we tested if iron chelators can compete with, and extract iron from, BTSL2-N and BTS-N. Iron release was followed in real time using colorimetric iron chelators such as Ferene^30^. The as-isolated, predominantly oxidized form of BTSL2-N displayed a slow but steady rate of approximately 20% iron release per hour when the protein was purified with β-mercaptoethanol in the buffers, or virtually no iron release without β-mercaptoethanol (Fig. 6a). However, if iron was released in the ferric form, it would not have been detected because Ferene specifically binds Fe^2+^. To investigate this possibility, we mixed β-mercaptoethanol-free BTSL2-N with the ferric iron chelator ethylenediaminetetraacetic acid (EDTA) and saw no change in the protein’s CD spectrum after 1 h (Supplementary Fig. 6). Thus, we conclude that, in their oxidized, Fe^3+^ form, the diiron sites are not labile.

**Fig. 6.**
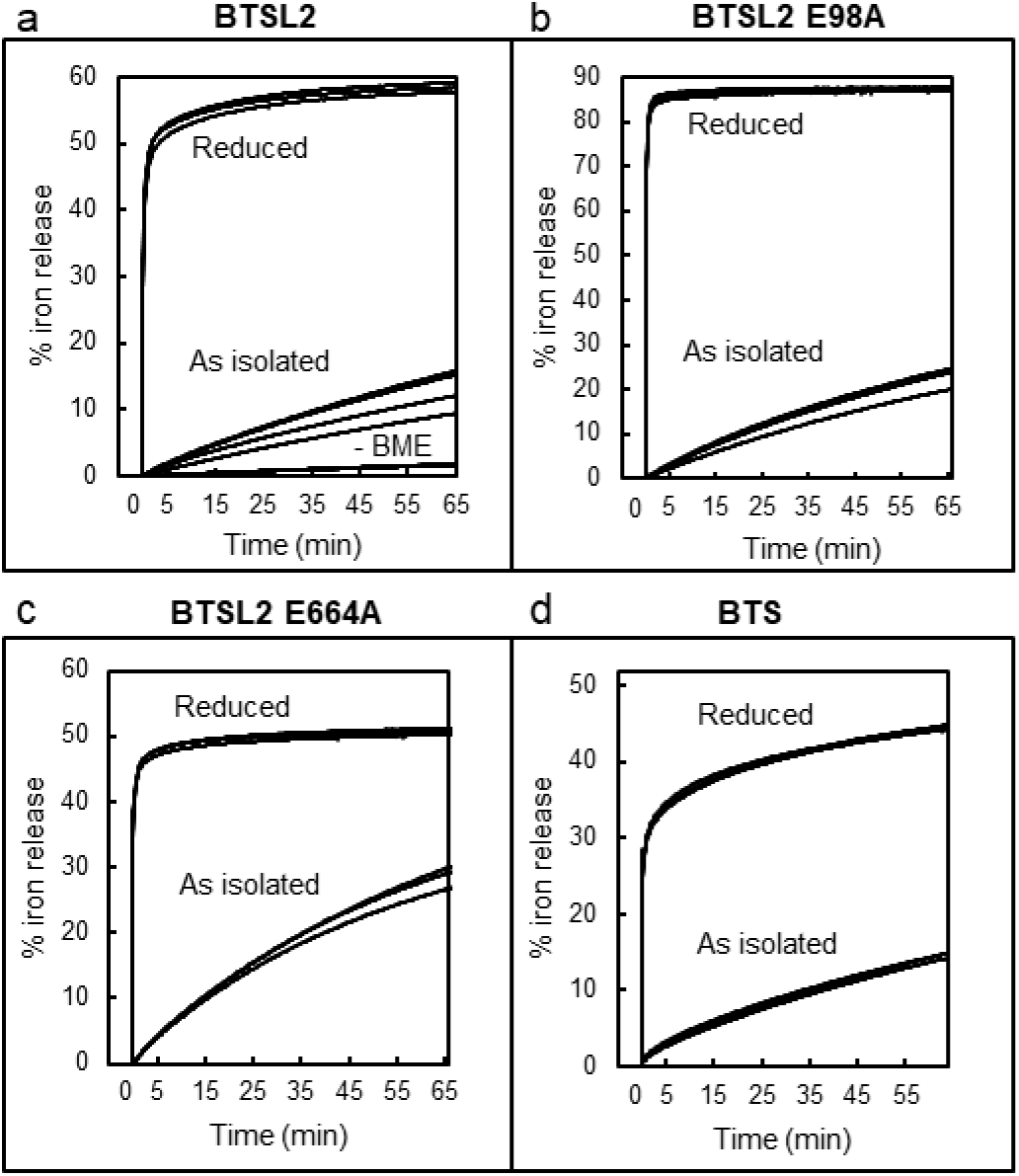
Iron binding to the Hr-like domain depends upon oxidation state. a-d. Release of protein-bound iron monitored using the colorimetric iron chelator Ferene. An increase in absorbance over time was correlated with the amount of iron released from the protein. Traces of independent time course measurements are overlaid. The ‘as isolated’ proteins are in the predominantly oxidized form but contain residual β-mercaptoethanol (BME) included to stabilize BTS. For BTSL2-N, protein isolated without BME is also shown. Proteins were reduced following incubation with dithionite and desalting. (**a**) BTSL2-N; (**b**) BTSL2-N E98A; (**c**) BTSL2-N E664A; (**d**) BTS-N.

Following reduction of BTSL2-N with dithionite and removal of excess reductant, exposure to Ferene resulted in release of ∼50% of the iron in the first few seconds, and then another 5% more slowly over an hour. For BTS-N, the oxidized sample released ∼15% iron over an hour, whereas the reduced protein rapidly released ∼30% and then another 20% at a much slower rate (Fig. 6d). Equivalent experiments with reduced BTSL2-N using two other colorimetric iron chelators, dipyridyl or ferrozine^31,32^, gave similar results in terms of the percentage of iron released and the kinetics (Supplementary Fig. 7). This demonstrated that the Fe^2+^-protein dissociation rate, rather than Fe^2+^-chelator association rate, was measured in these experiments.

For both BTSL2-N and BTS-N, not all of the iron was released, indicating that iron in one or more of the diiron centres is not labile, even when reduced. To further investigate this for BTSL2-N, iron release experiments were repeated with the E98A and E664A variant proteins. Reduced E98A BTSL2-N, in which the H3 diiron centre remains intact, released ∼85% of the iron in the first rapid phase but did not exhibit the slower release phase observed in the wild-type form (Fig. 6b). The reduced E664A protein, in which the H1 diiron centre remains intact, released ∼45% of its iron, mostly in the rapid first phase (Fig. 6c). Combining these values equates to a release of 67% iron, which is relatively close to the 60% released from BTSL2-N with both diiron centres. The data indicate that the H3 diiron centre is more labile, releasing up to both irons. The H1 diiron centre is also labile but releases one iron only. In addition, kinetics of both single variants appeared altered relative to the wild-type protein, suggesting that the iron status of one diiron centre affects the other.

To investigate whether iron binding is reversible, we tested for re-association of iron after the removal of the labile iron atoms. The experimental procedure, as outlined in Fig. 7a, was followed using CD spectroscopy because only protein-bound iron contributes to the spectrum. Under anaerobic conditions, a defined amount of Ferene was added to reduced BTSL2-N to remove 40% of the Fe^2+^ (i.e. less than the proportion removed in the fast phase when excess chelator was added, see Fig. 6a). To verify that the iron was indeed removed, half the sample was desalted and oxidized, yielding a CD spectrum with a commensurate decrease in the CD signal at 350 nm (Fig. 7b, blue trace). To the other half of the sample, kept under anaerobic conditions, a 3-fold excess of Fe^2+^ in the form of ferrous ammonium sulfate was added followed by a desalting step to remove unbound iron and iron-chelator complexes. After exposure to O_2_, the resulting CD spectrum was identical to that of the as-isolated protein (Fig. 7b, pink trace), indicating full occupancy of the diiron centres and successful re-association of the iron that had been removed. Overall, these data demonstrate that, in a reducing environment, iron binding to BTS/Ls is reversible. By contrast, in an oxidizing environment, iron remained tightly bound.

**Fig. 7.**
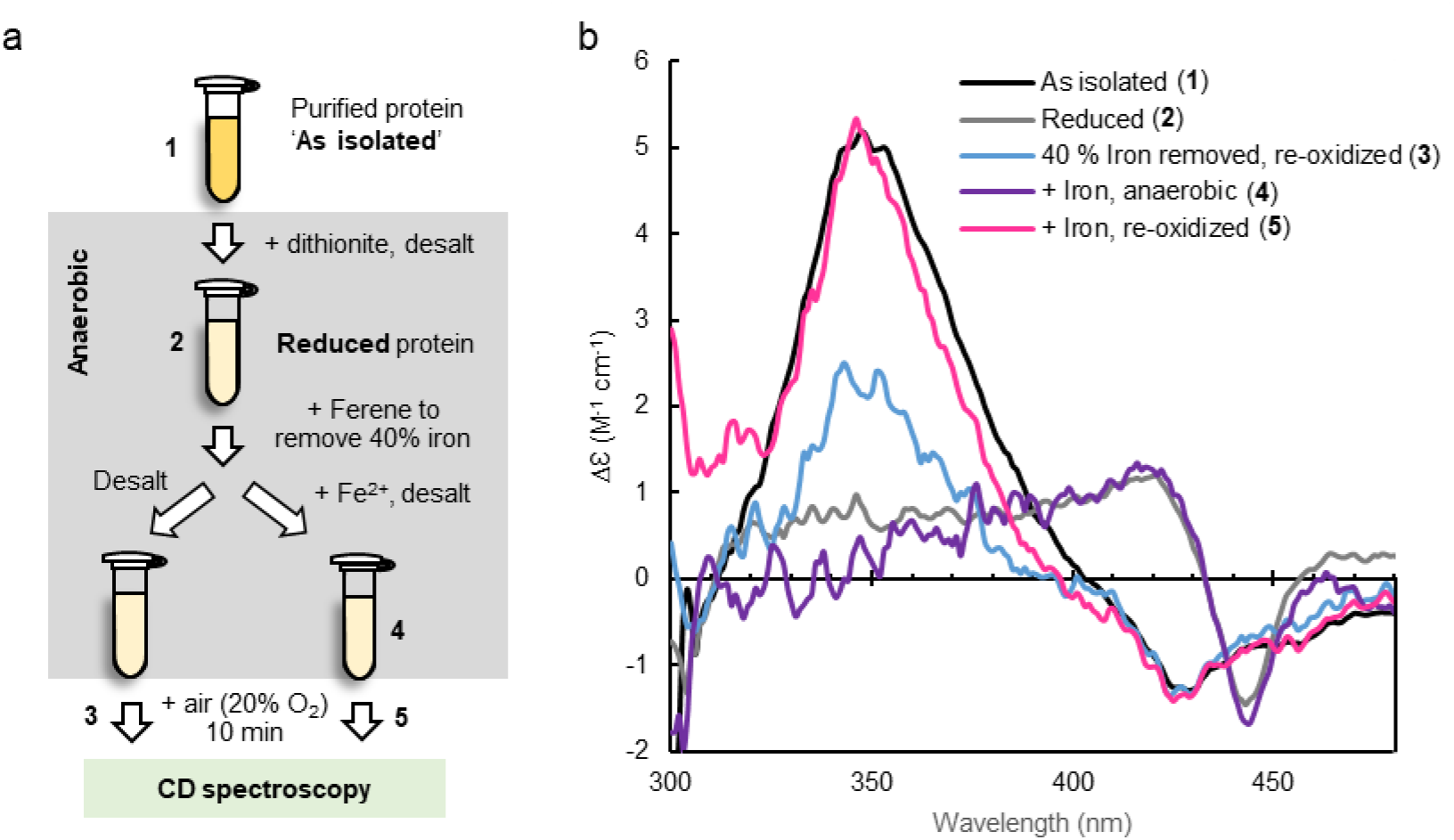
BTSL2-N can be fully reconstituted after partial iron removal. **a** Diagram of the experimental procedures to first remove labile iron and then reconstitute the diiron centres monitored by CD spectroscopy, which only detects protein-bound iron. **b** CD spectra of purified MBP-BTSL2-N (86 µM) as isolated (black) and after reduction with dithionite and its subsequent removal by desalting (grey). The reduced protein was mixed with a defined amount of Ferene, calculated to remove 40% of the protein-bound iron. Half the sample was desalted to remove Fe^2+^ chelator species, and exposed to O2 (light blue) to demonstrate the partial removal of protein-bound iron. The other half was mixed with a 3-fold excess of ferrous ammonium sulfate and desalted to remove unbound iron, keeping the protein reduced (purple), followed by exposure to O2 (magenta). Spectra were adjusted to compensate for dilution by reagent additions and desalting. The results are representative of two independent experiments.

### Impaired iron binding and redox activity in the H1 subdomain affects the function of BTS in vivo

We previously identified a mutation in the N-terminal domain of BTS that causes the plant to hyper-accumulate iron^33^. The in-frame deletion removes 5 amino acid residues from the first Hr-like sequence close to two iron-binding ligands (Fig. 8a). The phenotypic consequences of the mutation, called *dgl*, have been studied extensively in pea (*Pisum sativum*) since it was identified several decades ago^34,35^. The transcript levels of two iron uptake genes, *Ferric Reduction Oxidase 1* (*PsFRO1*) and *Iron-Regulated Transporter 1* (*PsIRT1*), were shown to be highly induced in *dgl* roots and did not decrease in response to addition of iron^36^. By contrast in wild-type plants, the levels of these transcripts were below the limit of detection under normal (+Fe) conditions. *FRO* and *IRT1* are downstream of the transcription factors ILR3/bHLH105 and bHLH115, which have recently been confirmed as ubiquitination substrates of BTS in Arabidopsis^2,5,9^. The Arabidopsis and pea *BTS* genes are functional orthologs^33^ and thus it can be deduced that the E3 ligase activity is diminished in the BTS*^dgl^*protein, upregulating the downstream signalling pathway and iron uptake genes.

**Fig. 8.**
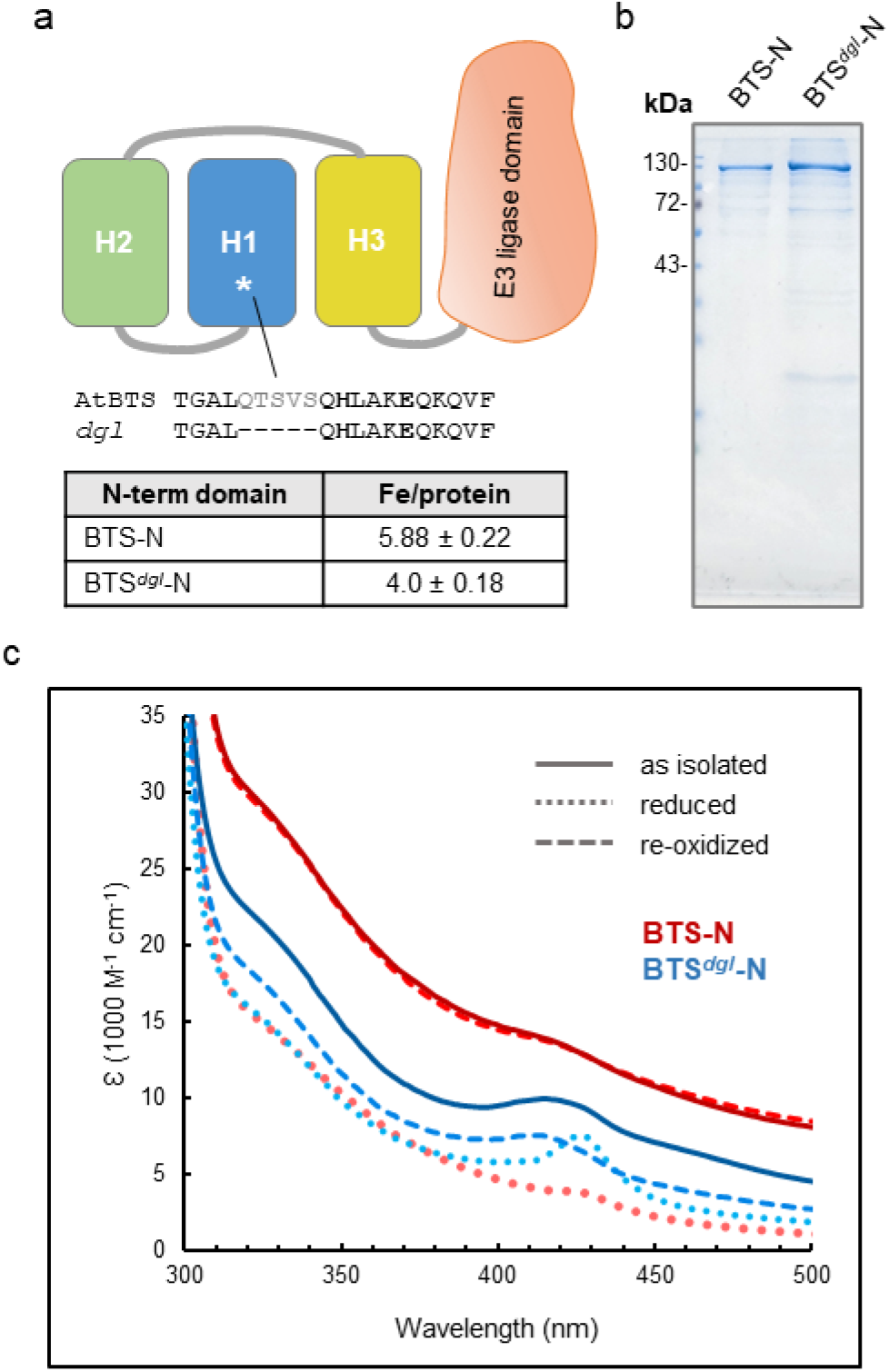
Lack of iron binding to H1 affects oxidation of the diiron centres and activity of BTS in vivo. **a** Diagram of the position of the 5 amino acid deletion in the *dgl* variant protein of BTS. The mutation identified in the *BTS* homologue in pea^33^, causing mis-regulation of iron homeostasis, was reconstructed in the Arabidopsis (At) expression construct of the N-terminal domain (BTS-N). The deletion is adjacent to two of the iron-binding ligands, H173 and E177, shown in bold. **b** Coomassie-stained SDS-PAGE gel of purified BTS-N (2.5 µg) and the *dgl* variant protein BTS*^dgl^*-N (5 µg), expressed as MBP fusion proteins in *E. coli*. The experimentally determined iron:protein ratio was 4 ± 0.18 for BTS*^dgl^*-N. **c** Redox reactions monitored by UV-visible spectroscopy of BTS*^dgl^*-N (blue-shaded lines) and BTS-N (red-shaded lines, from Fig. 3d) for comparison. The as-isolated proteins (solid lines) were reduced with dithionite (dotted lines) and re-oxidized by exposure to air for 50 min (dashed lines). Iron:protein rations and spectra of BTS*^dgl^*-N are representative of two independent protein purifications.

To investigate whether iron binding and/or redox activity is affected in the BTS*^dgl^* protein, the equivalent five amino acids were deleted from the Arabidopsis sequence in the BTS-N expression construct. The *dgl* variant protein was expressed and purified using the same conditions as for BTS-N (Fig. 8b). Although the yield of BTS*^dgl^*-N was ∼7-fold lower than for the wild-type protein, the mutant variant was equally stable through subsequent analyses. Iron quantification showed that the *dgl* variant bound 4 ± 0.18 iron per polypeptide, correlating with the lower molar absorptivity in the 300 – 500 nm range of the as-isolated protein compared to BTS-N (Fig. 8c, solid lines). The data indicated that the recombinant *dgl* protein lacked the diiron centre in the H1 subdomain.

Upon reduction with dithionite, the BTS*^dgl^*-N and BTS-N wild-type proteins showed similar UV-visible spectra (Fig. 8c, dotted lines), apart from the absorbance feature at 425 nm which was more pronounced in the *dgl* variant protein. Upon reoxidation by exposure to air, the UV-visible spectrum of wild-type BTS reverted to the as-isolated spectrum. However, for BTS*^dgl^*-N, only half the original absorptivity was recovered, even after longer incubation (50 min). Comparison of the EPR spectra showed that the minority mixed-valent signal was lacking in BTS*^dgl^* (Supplementary Fig. 8), indicating that this signal arises from or is dependent on the H1 diiron centre. Thus, impaired iron binding and reoxidation of BTS*^dgl^* in vitro are correlated with compromised functionality of the E3 ubiquitin ligase in vivo.

## DISCUSSION

Physiological and genetic studies firmly point to BTS/L proteins as the iron sensors that switch off the iron deficiency response in plants, but biochemical evidence has been lacking. Our in-depth study of the iron-binding and redox properties of the N-terminal Hr-like domain leads us to propose a mechanism by which BTS/L proteins are activated in response to increasing iron levels and oxidative stress (Fig. 9). The expression of BTS/L is induced by iron deficiency, so when the proteins are initially produced it is likely that the iron-binding sites are only partially occupied. Partial iron binding could be important for maintaining the protein in a folded, but inactive, state. As iron levels increase, the diiron centres fill. As and when the cellular environment experiences oxidative stress conditions, rapid oxidation of the diiron centres decreases the lability of the iron and shifts the protein into its active conformation, which is now able to ubiquitinate its cognate transcription factors, shutting down iron uptake.

**Fig. 9.**
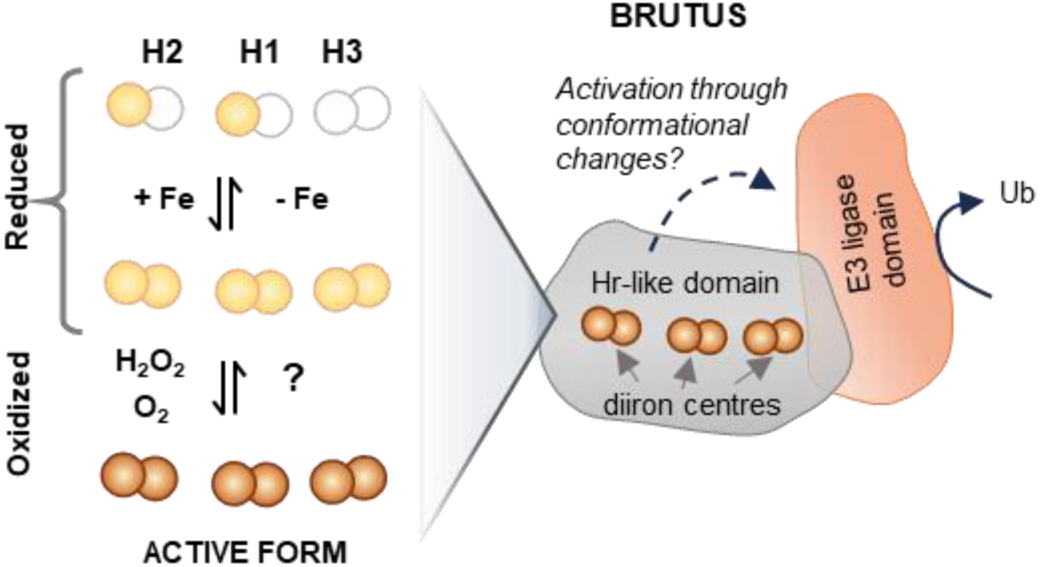
Proposed mechanism by which BTS/Ls are activated by iron binding and oxidation. In a reducing cell environment, iron binding is dynamic, with the diiron centres becoming fully occupied as the iron concentration increases. Oxidation of the diiron centres increases the binding affinity of iron, fixing the cofactors in place and leading to activation of the E3 ubiquitin ligase, presumably through conformational changes. Depicted here is the model for BTS with three diiron centres; the H2 diiron centre is missing in BTSL2. Fe^2+^ and Fe^3+^ are indicated by light brown and darker brown spheres, respectively.

The model makes certain assumptions and also leaves several open questions, to be tested in future experiments. First, the proposed mechanism would allow distinction between iron-induced oxidative stress and oxidative stress from other sources. We suggest that iron binding to BTS/L (without oxidation) is insufficient to trigger activation. As multicellular organisms, plants have cells that experience relatively high intracellular iron concentrations, for example vascular tissues involved in iron transport. Their role is to pass the iron on to neighbouring cells, and this flux should not be suppressed too easily by BTS/L activation. Having said that, unicellular algae also have BTS proteins, perhaps to maximize iron flow to chloroplasts which are a large sink of iron in photosynthetic cells.

Second, we speculate that the E3 ligase is activated through conformational changes that enable binding of the cognate E2 partner and/or substrate proteins, although other mechanisms are also possible. The observation that expression of BTS lacking the N-terminal triple Hr-like domain is more abundant and more active in planta^5^ suggests that this form is constitutively active. Possibly, the mostly rigid triple Hr-like domain, which is connected by a flexible linker to the E3 ligase domain, could have an auto-inhibitory function through large conformational changes. To investigate this, expression and purification of sufficient amounts of the full-length proteins is needed and efforts towards this are currently ongoing.

Interesting parallels between mammalian FBXL5 and plant BTS/L proteins are the use of diiron centres in Hr-like folds for sensing iron, linked to ubiquitin ligase activation. However, our characterization of the BTS/L Hr-like domains also revealed fundamental differences, which are likely to impact on the mechanism of iron sensing. Phylogenetic analysis (Fig. 1a) showed the invariant presence of three Hr-like subdomains in BTS/L, intertwined in a rigid triple domain (SAXS data, Fig. 2 and Supplementary Fig. 1). Several lines of evidence indicate that the three Hr-like subdomains of BTS/L do not function independently of each other. Most strikingly, the remaining intact diiron centres (H2 and H3) of the *dgl* variant BTS protein exhibited incomplete reduction and re-oxidation behaviour (Fig. 8), demonstrating that the absence of the H1 diiron centre affects the properties of the other two. This could be an effect on redox activity, or on iron binding affinity, or both. Inter-dependency of, or cooperativity between, the Hr motifs is consistent with the spectra (colour) of the two BTSL2 variants E98A and E664A (Fig. 3), which each lack one of the two diiron centres of the wild-type protein. As isolated, these variants exhibited absorption properties that were similar to each other but quite different form the wild-type protein, in that they lacked the absorbance shoulder at 350 nm. Independent behaviour would be expected to result in spectra similar to that of wild-type protein but of weaker intensity. Different behaviour was also noted in iron-release experiments, where the slow phase of release was much reduced in both variants (Fig. 6). Thus, while further data on the transduction of the signal from the Hr-like domain to the E3 ligase domain are needed, it is highly likely that the three Hr-like motifs function in a coordinated and inter-dependent manner that affects their redox properties.

Aside from the higher order architecture, the properties of the diiron centres appear similar in BTS/L proteins and FBXL5: The UV-visible spectra of BTSL2 (Fig. 3d) and FBXL5^37,38^ are comparable and both proteins have a relatively fast autoxidation rate (Fig. 4 and Refs. ^39–41^). AlphaFold models of BTS/Ls compared with the crystal structure of FBXL5^37^ suggest a similar coordination geometry of the diiron centres. The ligand binding is asymmetric, with one iron hexa- and the other iron penta-coordinated, which is likely to cause their differential lability as observed here for BTS/L proteins (Fig. 6b-c).

Iron binding stabilizes the FBXL5 protein^10,11^ and recent data showed that BTS from soybean is stabilized by iron when transiently expressed in tobacco (*N. benthamiana*) leaves^42^. However, previously a destabilizing effect by iron was reported when Arabidopsis BTS was expressed in a wheat-germ expression system^5^. We have repeated this experiment and found that iron inhibits the transcription or translation in this cell-free system but not the stability of the protein (Supplementary Fig. 9a). Attempts to remove all iron from oxidized BTSL2-N gradually by increasing the temperature in the presence of an iron chelator resulted in precipitation of the protein (Supplementary Fig. 9b). From these experiments and our iron binding studies (Fig. 6 and 7), we conclude that the BTS/L N-terminus with partially occupied diiron centres is folded, and that iron is required for stability of the protein.

The need for a thiolate reductant such as β-mercaptoethanol or glutathione to purify stable BTS-N suggests that glutathione plays a physiological role in the iron sensing mechanism. Decreased glutathione levels as a consequence of mutations in the biosynthesis pathway render Arabidopsis seedlings more sensitive to iron deficiency. In those mutants, key genes required for iron uptake, including *IRT1* and *FRO2*, were expressed at much lower levels than in wild-type plants^43^. It is tempting to speculate that in these mutants the redox equilibrium of the BTS/L proteins is shifted more towards the oxidized form (Fig. 9), leading to increased ubiquitination followed by degradation of the transcription factors of the iron deficiency response.

The iron-binding and redox properties of the unique triple Hr-like domain of BTS/L proteins shed the first light on the sensing mechanism of these important negative regulators of the iron deficiency response in plants. This will facilitate mutant studies to probe the mechanism in planta, and to potentially modify the system for increasing iron uptake in specific plant tissues for human nutrition, known as biofortification.

## METHODS

### Protein expression and purification

Coding sequences of the N-terminal domains of *BTS* (AT3G18290) and *BTSL2* (AT1G18910) were amplified from *Arabidopsis thaliana* cDNA using the primers listed in Supplementary Table 3 and cloned into the pMAL-c5x vector (New England Biolabs). The variant BTSL2 and BTS proteins were made by site-direct mutagenesis using the listed primers and either the QuikChange Lightning Site-Directed Mutagenesis Kit (Agilent) or the QS Site Directed Mutagenesis Kit (New England Biolabs) according to the supplier’s manual. For the BTS*^dgl^* variant, the linearized plasmid excluding the 15-nucleotide deletion was blunt-end ligated.

The resulting plasmids were used to transform BL21-DE3 Rosetta-pISC *E. coli* cells^44^. The transformed cells were cultured in 4 L Terrific broth containing 0.02% (v/v) glucose at 37 °C to an OD_600_ of 1.1-1.2. Flasks were incubated in an ice bath for 15 min, then induced with 100 µM IPTG, 0.05% (v/v) DMSO and 50 µM ferric ammonium citrate o/n at 16 °C. Cells were harvested and resuspended in 100 mM Tris-HCl, 150 mM NaCl, 20 mM β-mercaptoethanol, pH 8.0 with cOmplete EDTA-free protease inhibitor (1 tablet/ 50 mL). Cells were disrupted using an ultrasonics processor (Vibra-Cell VCX 750, Sonics) during 60 cycles of 2 s on, 8 s off at 40% power while samples were cooled in ice water. The resulting lysate was centrifuged at 20,000 × g for 30 min at 4 °C. The soluble fraction was loaded onto a 5 mL Strep-Tactin XT column (IBA Lifesciences) and washed with 100 mM Tris-HCl, 150 mM NaCl, 20 mM β-mercaptoethanol, pH 8.0. Protein was eluted with 100 mM Tris-HCl, 150 mM NaCl, 20 mM β-mercaptoethanol, 50 mM D-biotin, pH 8.0. Removing β-mercaptoethanol from purification buffers after sonication for MBP-BTSL2-N had no effect on the final Fe:protein ratios. Proteins were concentrated using a spin concentrator (MWCO 30 kDa, Pierce). For SAXS experiments, affinity-purified MBP-BTSL2-N-Strep was cleaved using Factor Xa protease (NEB, product no. P9010L) at a concentration of ∼2.8 ng mL^-1^ for 24 h at 4 °C. Cleaved protein was buffer-exchanged into 10 mM MES, 15 mM NaCl, pH 6.5 using a spin concentrator (MWCO 30 kDa, Pierce) and BTSL2-N was isolated via size-exclusion chromatography using an S200 26/60 Sephadex column (Cytiva). The identity of the purified BTS-N and BTS*^dgl^*-N variant proteins was confirmed by LC-MS of the intact proteins.

### Quantification of iron and other metals

Protein samples were buffer exchanged into 10 mM Tris-HCl, pH 8.0 and analysed using inductively coupled plasma – mass spectrometry (ICP-MS; Triple Quadrupole iCAP-TQ from ThermoFisher). Iron in protein samples was also quantified using the colorimetric reagent 3-(2-pyridyl)-5,6-bis(2-(5-furyl sulfonic acid)-1,2,4-triazine, also called Ferene^30^. Protein samples were quantified using the Bradford assay (Protein Assay Dye Reagent, BioRad).

### Small-angle X-ray scattering

SAXS data were collected at Diamond Light Source (DLS; Oxfordshire, UK) on beamline 21 in a standard SEC-SAXS experiment. Samples at 1 to 5 mg mL^-1^ were applied to a Superdex 200 increase column (Cytiva, Amersham, UK) and run at a flow rate of 0.075 mL min^-1^ in 10 mM MES, 15 mM NaCl, pH 6.5. Data were recorded at a wavelength of 0.9464 Å using an EigerX 4M detector (Dectris, Baden-Daettwil, Switzerland). Measurements were performed at 15 °C within the *q* range 0.0045 – 0.34 Å^-1^ (*q* = 4πsinθ/λ; 2θ is the scattering angle, λ is the wavelength) with a sample-to-detector distance of 3.72 m. Data were normalized for water scattering on an absolute scale. From the SEC-SAXS run of the 5 mg mL^-1^ protein sample frames 372-375 were selected, merged and solvent subtracted for data analysis using ScÅtter (https://sibyls.als.lbl.gov/scatter/), IV.d and ATSAS 3.2.1^45^. Calculation of *R*_g_ and forward scattering *I*(0) was performed by Guinier approximation after removing the first 30 data points, The maximum dimension of the particle *d*_max_ and the *P(r)* function were estimated using the program GNOM^46^. Ab initio envelope models were computed using DAMMIN^19^ and overlaid with an AlphaFold2 model^47,48^ that was refined against the scattering data using SASREF^49^. The bound iron centres were populated manually via structure superposition with existing PDB entry 2AVK and model editing in the COOT software^50^. Experimental data and models were prepared using IUCr guidelines^51,52^, as summarized in Supplementary Data 1.

### Titrations with O_2_ or H_2_O_2_

The MBP-fusion proteins of the BTSL2 and BTS N-terminal domains were stable at room temperature in a glovebox over a 12 h period, during which redox studies could be performed in an anaerobic environment. Buffer solutions were sparged with N_2_ gas for 1 h before being left to equilibrate in an anaerobic (<2 ppm O_2_) chamber overnight. Protein aliquots were thawed inside the chamber. Gas-tight quartz cuvettes with a 10 mm pathlength (Starna Cells, UK) were used to transfer the protein in and out of the chamber for measurement. Sodium dithionite was prepared in anaerobic buffer and its concentration was determined using an extinction coefficient, ε_315_ = 8000 M^-1^ cm^-1^ (Ref^53^). A threefold excess of dithionite was added relative to the protein’s iron content and incubated anaerobically. Reduction of the protein was confirmed using circular dichroism measurements made on a Jasco J-810 CD spectropolarimeter to monitor loss of the protein’s absorbance peak due to Fe^3+^ at ≈350 nm. Protein was then buffer exchanged on a PD10 desalting column (Cytiva). Aliquots of protein were then titrated with known volumes of H_2_O_2_ prepared in anaerobic buffer or with aerobic buffer. After each addition of oxidant, a minimum of three minutes elapsed before recording the sample’s absorbance or CD spectrum. Absorbance spectra were recorded using a Jasco V550 spectrophotometer.

### Electron Paramagnetic Resonance

Proteins were degassed, reduced and desalted as above. Proteins were mixed within an anaerobic chamber with the appropriate volume of a supersaturated O_2_ solution (prepared by bubbling pure O_2_ through 100 mM Tris-HCl, 150 mM NaCl, pH 8.0 in a suba-sealed volumetric flask), estimated to be 1.2 mM, or a 1.2 mM H_2_O_2_ solution, and then transferred into EPR tubes. For titrations with sodium dithionite, aliquots of protein were mixed in an anaerobic chamber with the appropriate volume of a 1.5 mM dithionite solution in plastic cannulae and ejected into EPR tubes. Sealed tubes were removed from the chamber and frozen after the stated reaction times by dropping into methanol cooled in dry ice. Spectra were recorded on a Bruker E500 (X band) EPR spectrometer at 10 K equipped with an Oxford Instruments liquid-helium system. Quantification of the paramagnetic species was done by comparison of second integrals of the species’ pure line shape EPR signals, obtained by the spectra subtraction with variable coefficient procedure^54^, with those of the two concentration standards used. For quantification, the microwave power saturation dependences of all signals were measured and taken into account (Supplementary Fig. 10). The mixed valent (Fe^2+^/Fe^3+^) species was quantified using a 1 mM Cu^2+^[EDTA] concentration standard, and the mono-nuclear ferric signals using a 35 µM ferric iron binding protein from *Neisseria gonorrhoeae*.

### Amplex Red assay

H_2_O_2_ release from reduced BTS/L proteins was detected using an Amplex Red kit (Thermo Fisher) according to the manufacturer’s instructions. Briefly, a 200 µL aliquot of freshly prepared aerobic Amplex Red reaction mixture was mixed with a 200 µL anaerobic aliquot of reduced BTS/L protein, diluted so that the protein-bound iron concentration was 40 µM. The absorbance at 571 nm was used to quantify H_2_O_2_ based on a calibration curve using 0-37.5 µM H_2_O_2._

### Iron chelator competition assays

For ‘as-isolated’ measurements, proteins were thawed in an anaerobic glovebox. For the reduced samples, proteins were treated with dithionite and desalted as described above. Each protein sample was then diluted in a sealed cuvette and single wavelength time courses were recorded using a Jasco V550 spectrophotometer after mixing with colorimetric chelators. Monitored wavelengths were 592, 525, and 562 nm for Ferene, 4,4’ dipyridyl and ferrozine, respectively. The final concentration of iron in each assay was 75 µM, and that for each chelator was 1 mM. The percentage iron released was measured relative to calibration curves prepared for each chelator mixed with 0, 10, 25, 50 and 75 µM Mohr’s salt anaerobically prepared in 100 mM Tris-HCl, 150 mM NaCl, pH 8.0.

### Reconstitution of BTSL2

BTSL2-N (86 µM, 3.8 Fe per protein) was reduced and desalted as above. Ferene sufficient to remove 40% of the iron within the protein was added, and iron removal confirmed using absorbance at 593 nm. Ferrous ammonium sulfate in 3-fold excess over freshly de-occupied iron binding sites was added to the protein under anaerobic conditions, after which the sample was again desalted. The protein was exposed to air for 10 min in an open Eppendorf with gentle mixing. UV-visible absorbance and CD spectra were recorded after each manipulation. Each CD spectrum represents the average of four measurements to improve signal:noise ratio; for the final sample (air exposed, reconstituted BTSL2) the spectrum was the average of sixteen measurements.

## Supporting information

Supplementary Data 1 (SAXS data)

## Acknowledgements

We thank our colleagues: Dr. Myles Cheesman for use of the CD spectrometer; Graham Chilvers for ICP-MS; Jason Crack for protein mass spectrometry, technical advice and discussion; Dominic Campopiano (University of Edinburgh) for providing ferric iron binding protein from *Neisseria gonorrhoeae*; Mark Thornton, Mei Ying Nam and Robert Griffith (summer students) for protein expression. We acknowledge Diamond Light Source for access to SAXS beamline B21 under proposal MX25108 and beamline scientist Katsuaki Inoue for collecting the data. This work was supported by the Biotechnology and Biological Sciences Research Council, grant awards BB/N001079/1 (to J.R.C, M.F., N.E.L.B. and J.B.); BB/V015095/1 (to J.P., M.F. and J.B.); BB/V014625/1 (to J.M.B. and N.E.L.B.); and BB/R013578/1 for the purchase of an ICP-QQQ-MS instrument (J.B. and N.E.L.B). J.R.C. received additional support from the NextGenerationEU program, award TED2021-130539A-I00; and MCIN/AEI/10.13039/501100011033.

## Author contributions

N.E.L.B. and J.B. conceived the project. J.P. carried out the majority of the experiments with guidance from J.M.B. J.R.-C. and M.F. made the expression constructs. J.E.A.M. performed SAXS analysis. D.A.S. supervised the EPR analysis. J.B. wrote the paper and all authors reviewed, edited, and commented on the draft. N.E.L.B. and J.B. jointly supervised the project.

