## Supplementary Figures and Tables for "Iron-sensing and redox properties of the hemerythrin-like domains of Arabidopsis BRUTUS and BRUTUS-LIKE2 proteins"

| | |
|---|---|
| Supplementary Fig. 1 | Additional SAXS data for the N-terminal domain of BTSL2. |
| Supplementary Fig. 2 | Redox activity of the N-terminal domain of BTSL2 monitored by CD spectroscopy. |
| Supplementary Fig. 3 | BTSL2-N is slowly reduced by glutathione. |
| Supplementary Fig. 4 | Characterizaton of mixed valent diiron signals in BTSL2-N and BTS-N by EPR. |
| Supplementary Fig. 5 | EPR spectrum of the free radical observed in BTSL2-N. |
| Supplementary Fig. 6 | Ferric iron is not released from the Hr-like subdomains of BTSL2. |
| Supplementary Fig. 7 | Iron release kinetics of reduced BTSL2-N are independent of the iron chelator properties. |
| Supplementary Fig. 8 | The BTS$^{dgl}$ variant protein shows no mixed-valent diiron species. |
| Supplementary Fig. 9 | BTS/L proteins are stabilized by iron. |
| Supplementary Fig. 10 | Power saturation curves for the EPR signals observed in BTS/L-N proteins. |
| Supplementary Table 1 | BTS and BTSL protein sequences for phylogenetic analysis. |
| Supplementary Table 2 | Element analysis of purified BTS-N and BTSL2-N. |
| Supplementary Table 3 | Primers used in the preparation of recombinant protein expression constructs for this study. |

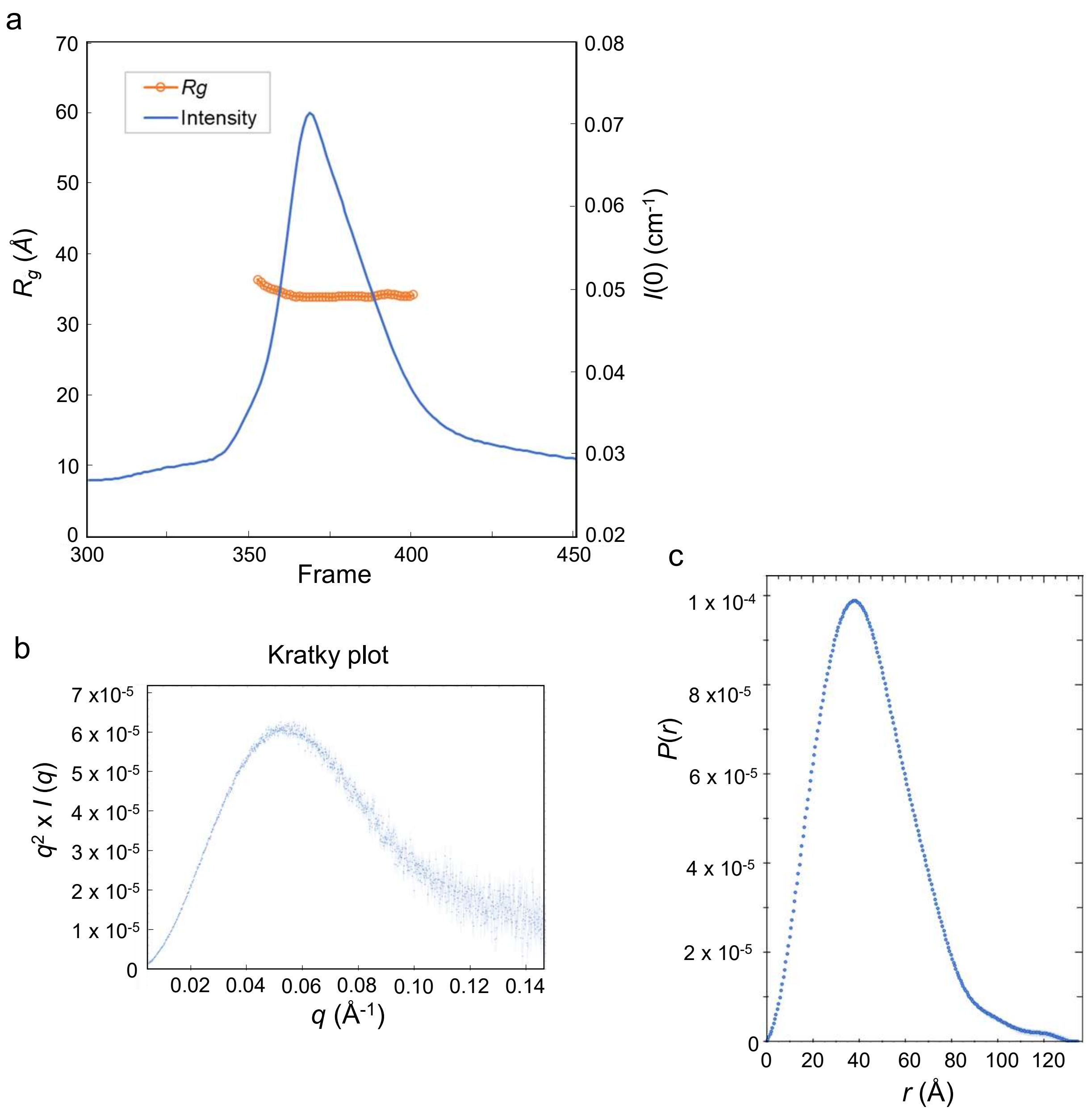


**Supplementary Fig. 1 - Additional SAXS data for the N-terminal domain of BTSL2. a** Elution profile of the N-terminal domain of BTSL2 (BTSL2-N), diluted to 5 mg mL$^{-1}$, monitored using scattering intensity (blue) in SEC-SAXS analysis. The radius of gyration $R_g$ (orange) was calculated for individual fractions (frames) of the main elution peak. **b** Kratky plot of $q^2$ x $I(q)$ versus $q$. **c** $P(r)$ versus $r$ profile (α = 1.007). Source data have been deposited under accession SASDU79 in www.sasbdb.org (www.sasbdb.org/data/SASDU79/1g65egyk1i/).

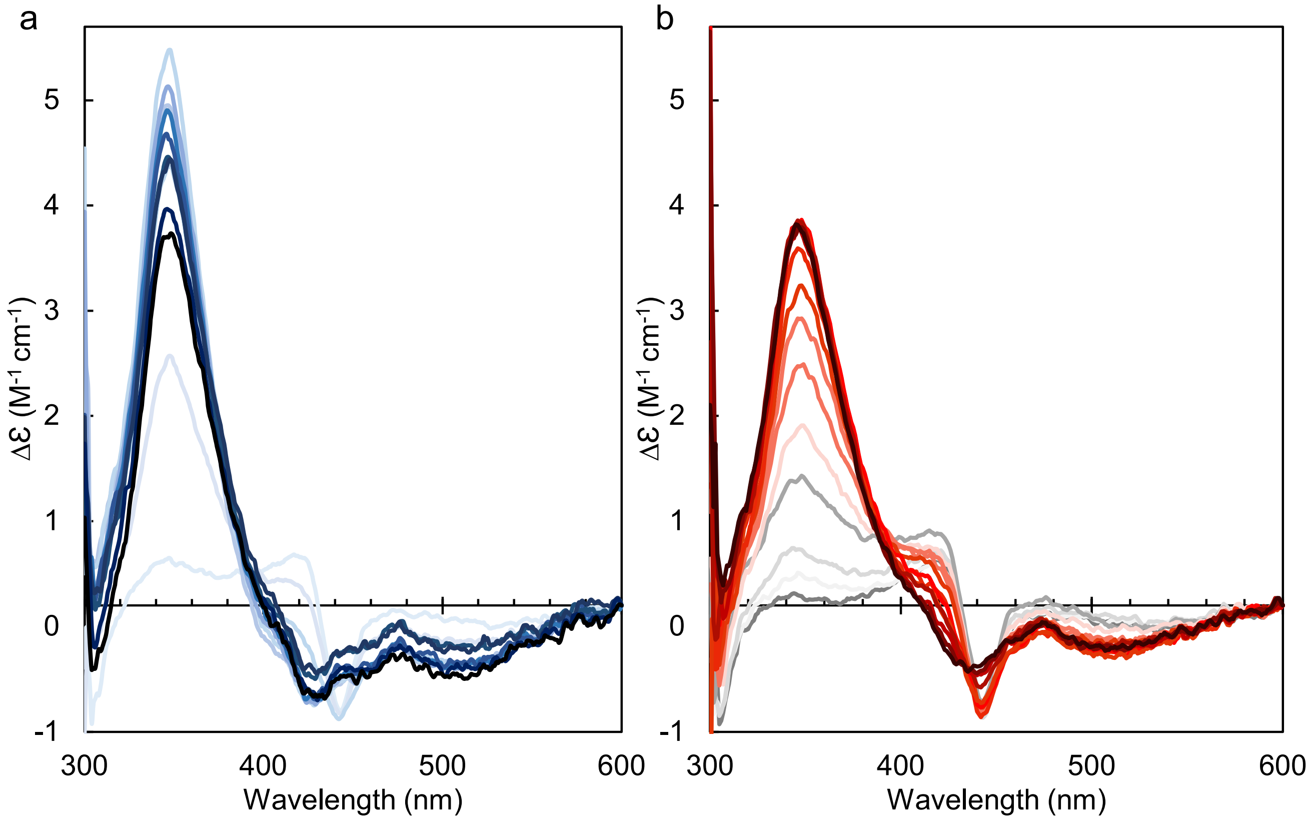


**Supplementary Fig. 2 - Redox activity of the N-terminal domain of BTSL2 monitored by CD spectroscopy. a-b** CD spectra of MBP-BTSL2-N (140 µM) monitoring titration with $O_2$ (a) or $H_2O_2$ (b). Black line: 'as-isolated' protein purified in air (a) or the end point of the titration (b). Grey line: spectrum after reduction with dithionite and desalting in an anaerobic chamber. Spectra were adjusted to compensate for dilution by reagent additions and desalting.

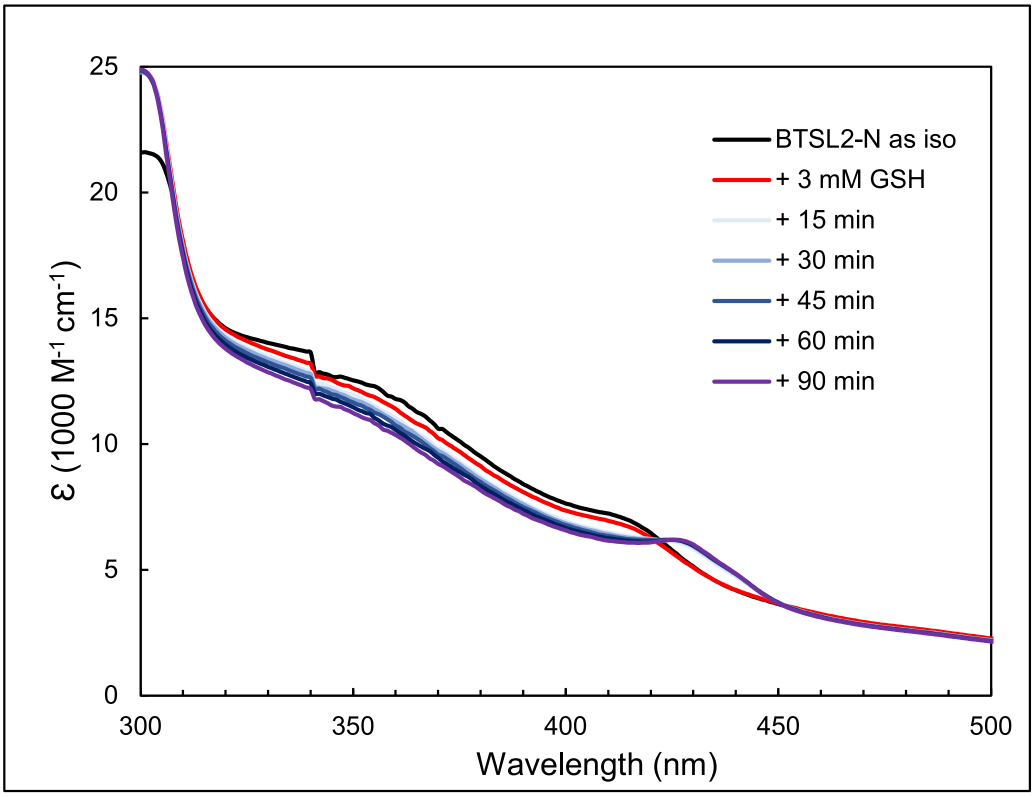


**Supplementary Fig. 3 - BTSL2-N is slowly reduced by glutathione.** UV-visible absorbance spectra of BTSL2-N (173 µM) 'as isolated' (black line); after transfer to an anaerobic environment for 30 min before being mixed with 3 mM glutathione (GSH, red line). Spectra were recorded after 15, 30, 45, 60 and 90 min.

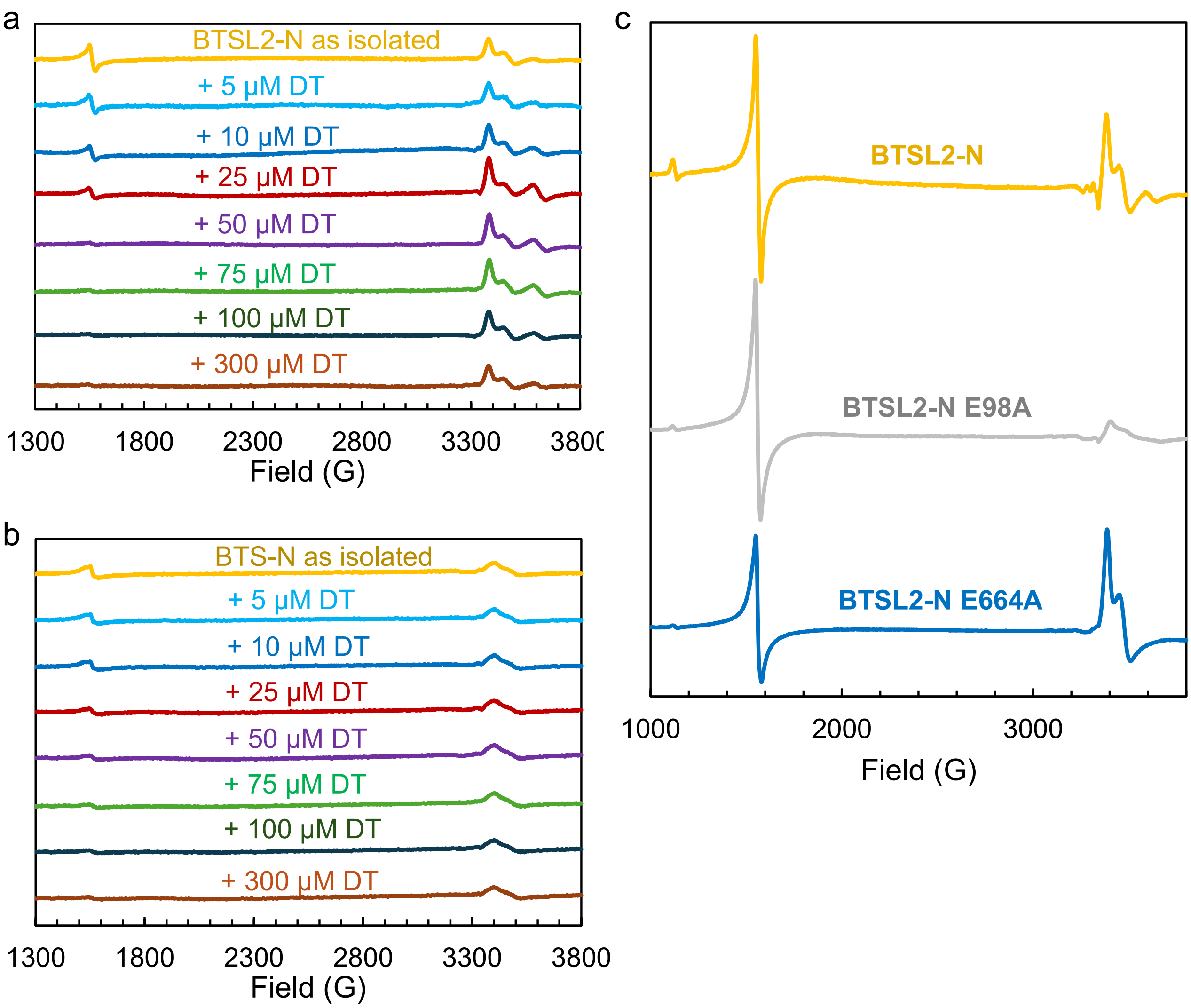


**Supplementary Fig. 4 - Characterization of mixed valent diiron signals in BTSL-N and BTS-N by EPR.**

**a-b** EPR spectra of anaerobic titrations with dithionite of (a) BTSL2-N (50 µM, 200 µM Fe) and (b) BTS-N (33 µM, 200 µM Fe). Protein samples were incubated for a minimum of 30 minutes in an anaerobic chamber, to exclude oxygen, and diluted with anaerobic buffer to a concentration of 200 µM (i.e. 50 µM for BTSL2-N and 33 µM for BTS-N). Aliquots of protein were then mixed with increasing amounts of dithionite, transferred to EPR tubes, capped, and removed from the chamber to be frozen in methanol, cooled by dry ice. The reaction time was 2 min from adding dithionite to freezing. **c** EPR spectra of WT BTSL2-N (142 µM, 3.9 Fe/protein) and E98A (345 µM, 1.9 Fe/protein) and E664A (527 µM, 2 Fe/protein) variants, as isolated.

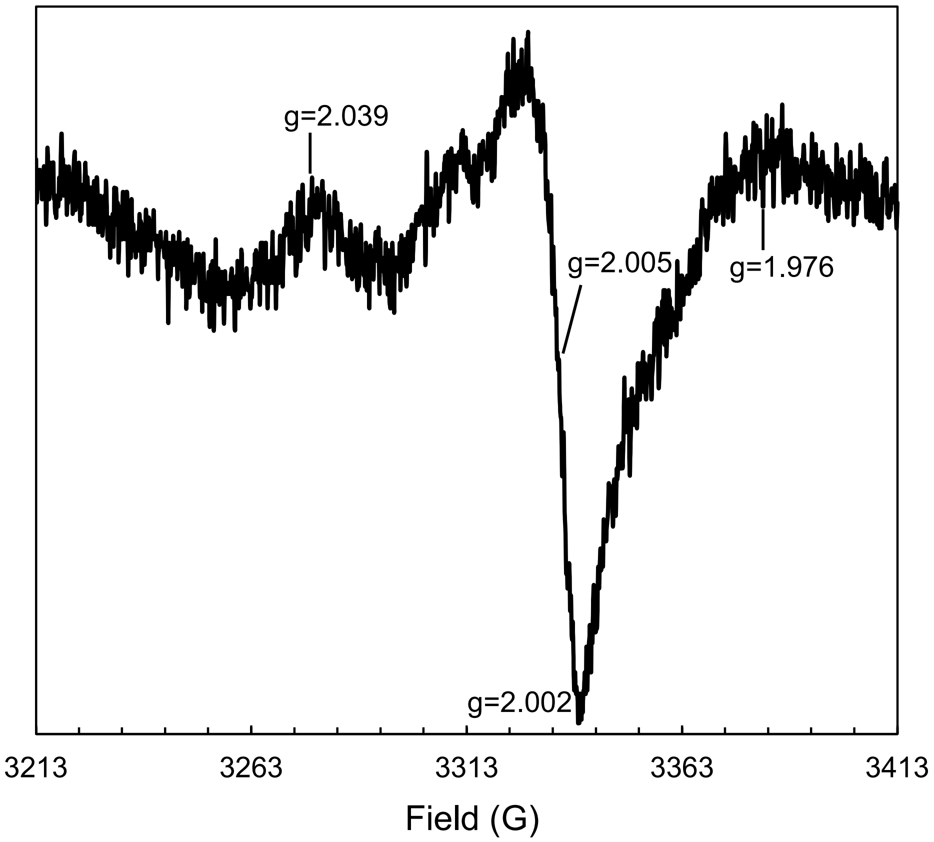


**Supplementary Fig. 5 - EPR spectrum of the free radical observed in BTSL2-N.** MBP-BTSL2-N (287 µM) as isolated with g-values of the major signals indicated. EPR spectra were recorded at 10 K using the following parameters: microwave frequency $v_{MW}$ = 9.353 GHz, modulation frequency $v_M$ = 100 kHz, time constant T = 82 ms, microwave power $P_{MW}$ = 3.17 mW, modulation amplitude $A_M$ = 5 G, scan rate v = 22.6 G $s^{-1}$.

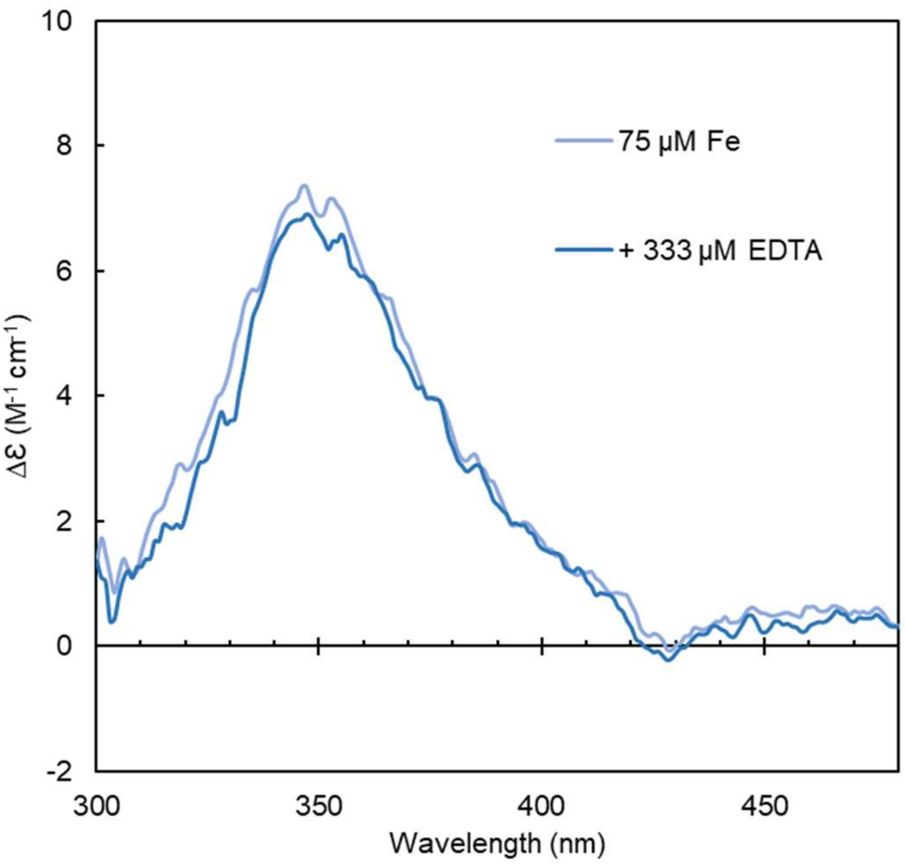


**Supplementary Fig. 6 - Ferric iron is not released from the Hr-like subdomains of BTSL2.** CD spectrum of the N-terminal Hr-like domain of BTSL2 'as isolated' (19 μM, 75 μM Fe), mixed under air with 333 μM EDTA and incubated at room temperature for 1 h before a second spectrum was recorded.

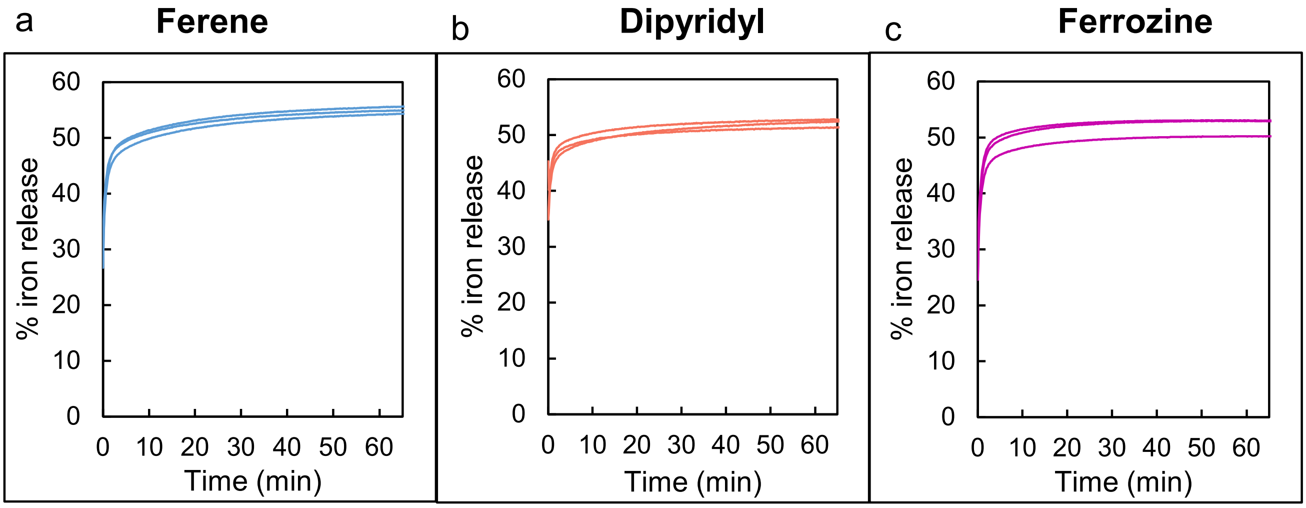


**Supplementary Fig. 7 - Iron release kinetics of reduced BTSL2-N are independent of the iron chelator properties.** Reduced MBP-BTSL2-N (19.7 µM protein, 75 µM iron in the 'as isolated' protein) was mixed with an excess of iron chelator (1 mM) Ferene (a), dipyridyl (b), or ferrozine (c). The increase in absorbance was measured over time to calculate the percentage iron release.

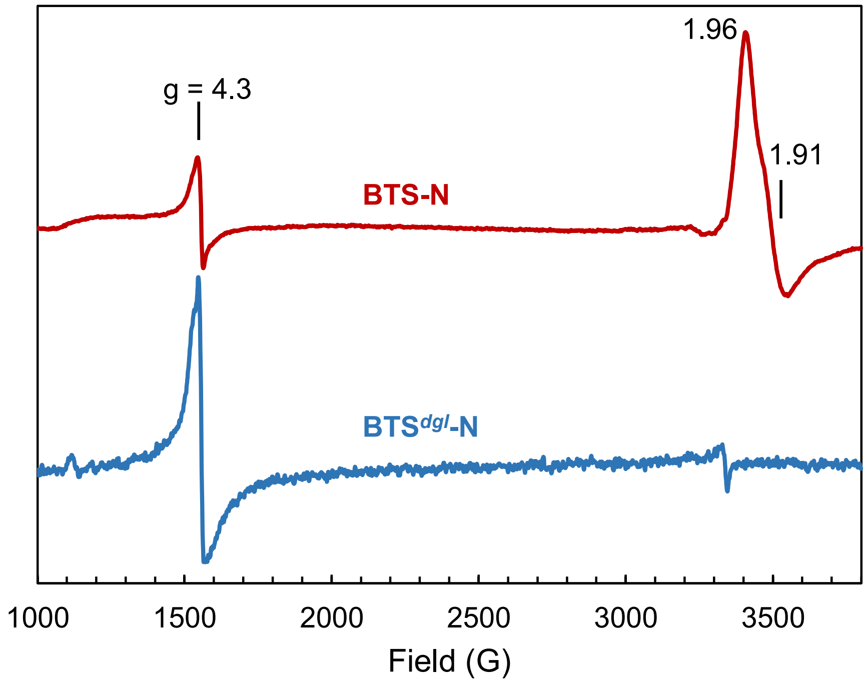


**Supplementary Fig. 8 - The BTS*dgl* variant protein shows no mixed-valent diiron species.** EPR spectra of BTS-N (222 µM, 5.8 Fe/protein) and BTS*dgl*-N (56 µM, 3.8 Fe/ protein), as isolated. The BTS-N spectrum is the same as shown in Fig. 5. The mixed-valent signal has g values of 1.96 and 1.91. Spectra have been scaled relative to their concentration.

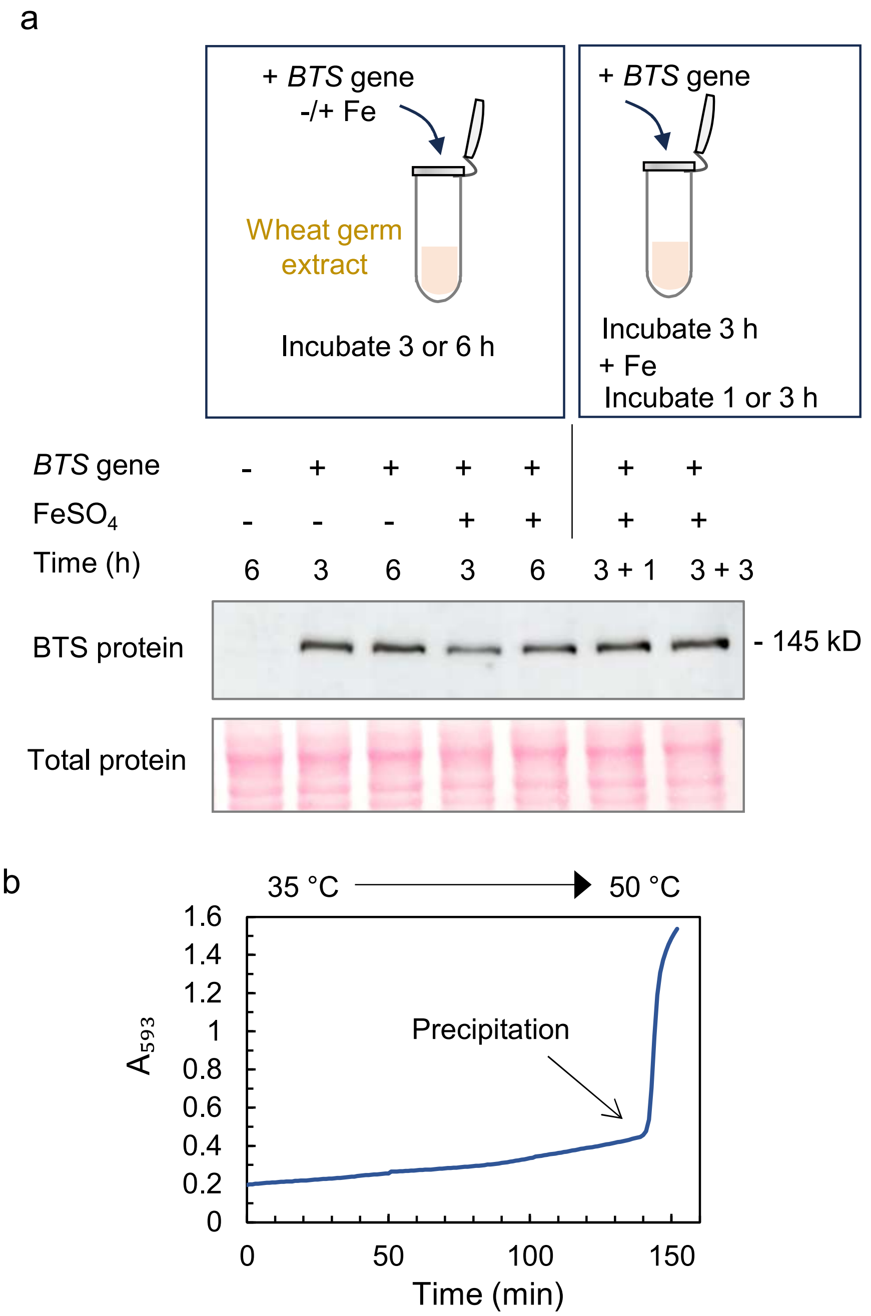


**Supplementary Fig. 9 - BTS/L proteins are stabilized by iron.**
**a** Production of BTS protein is inhibited by iron in a wheat germ expression system. Repeating a previously reported experiment[1], we used the Arabidopsis *BTS* coding sequence as template in the TnT SP6 High-Yield Wheat Germ Protein Expression System (Promega), closely following the supplier's manual. Ferrous sulfate, 0.5 mM final concentration, was added at the start or after 3 hours. Aliquots taken at the indicated time points were analysed by Western blot analysis with antibodies against BTS kindly donated by Terry Long, North Carolina Sate University, and the same as in Ref. 1). Total protein per lane was visualised by Ponceau S staining of the membrane. **b** Physical removal of bound iron from the BTSL1 N-terminal domain (BTSL1-N) leads to precipitation. BTSL1-N (12 µM, 3.3 Fe/protein) was incubated with 100 µM ascorbate and 200 µM Ferene. The sample was incubated at 35 °C, increasing the temperature by 5 °C every 16 min until the protein aggregated at 50 °C.

[1]Selote, D., Samira, R., Matthiadis, A., Gillikin, J.W. & Long, T.A. Iron-binding E3 ligase mediates iron response in plants by targeting basic helix-loop-helix transcription factors. *Plant Physiol.* **167**, 273-86 (2015).

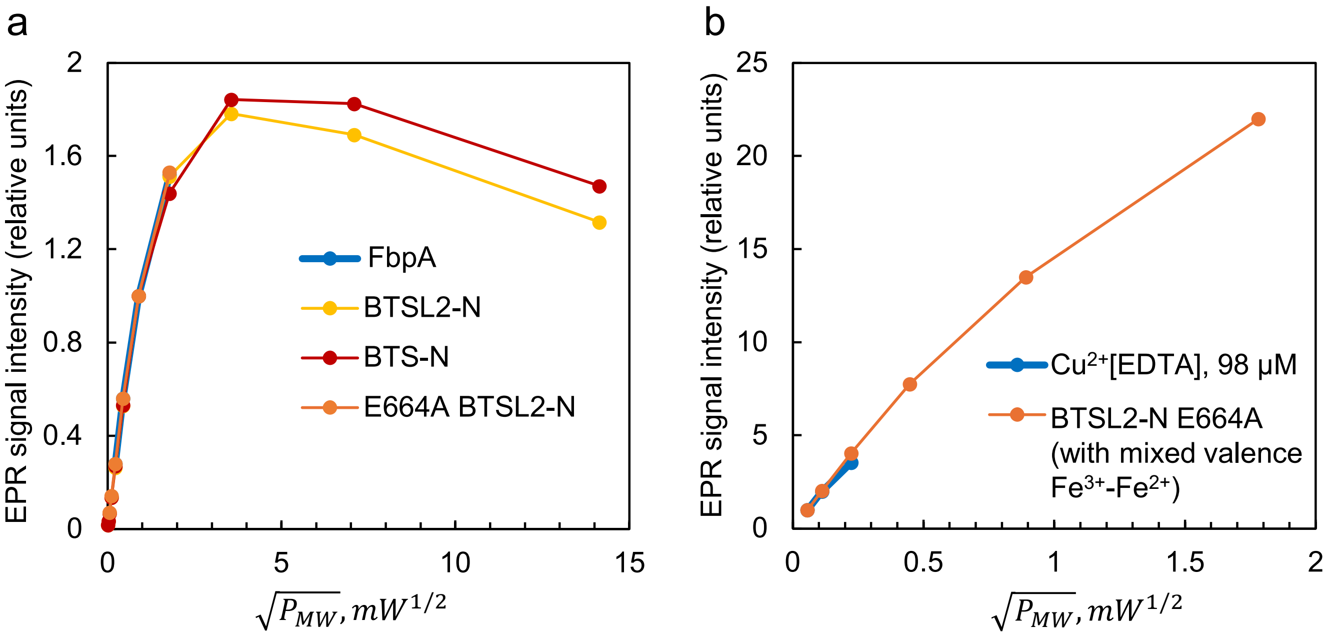


**Supplementary Fig. 10 - Power saturation curves for the EPR signals observed in BTS/L-N proteins. a** Power saturation curves for the g = 4.3 signals of Ferric iron binding protein FbpA (blue), BTSL2-N (yellow), BTS-N (red) and BTSL2-N E664A (orange). Each curve is normalised to an identical slope for the initial linear parts of the dependences (no saturation at low powers) with intensity 1.00 set for the microwave power of $P_{MW}$ = 0.8 mW.

**b** Power saturation curves for a 98 μM $Cu^{2+}$[EDTA] standard, and for the EPR signal from the mixed-valent state of the diiron centre in BTSL2-N E664A (527 μM). The two curves are normalised as in (A) with intensity 1.00 set for the microwave power of $P_{MW}$ = 0.00317 mW.

**Supplementary Table 1. BTS and BTSL protein sequences for phylogenetic analysis**.

| **Common name** | **Scientific name** | **Classification** | **UniProt Entry**<br>or Phytozome v13 gene |
|---|---|---|---|
| Rice | *Oryza sativa* | Monocots, Poaceae | V9G2Z0<br>B8AWE7 |
| Maize | *Zea mays* | Monocots, Poaceae | A0A1D6NF39<br>A0A1D6MBU0 |
| Barley | *Hordeum vulgare* | Monocots, Poaceae | F2DJA0<br>A0A8I6WU11 |
| Wheat | *Triticum aestivum* | Monocots, Poaceae | A0A3B6EJS9<br>A0A3B5Y5G3 |
| Vanilla | *Vanilla planifolia* | Monocots, Orchidaceae | A0A835V099 |
| Arabidopsis (thale cress) | *Arabidopsis thaliana* | Dicots, Brassicaceae | Q8LPQ5<br>F4IDY5<br>F4HVS0 |
| Medicago (barrel medic) | *Medicago truncatula* | Dicots, Fabiaceae | G7L9R6<br>A0A072V1D4 |
| Spinach | *Espinachia oleracea* | Dicots, Amaranthaceae | A0A9R0I415<br>A0A9R0KD67 |
| Water lily | *Nymphea colorata* | Basal angiosperms, Nymphaeaceae | Nycol.G00048 |
| Amborella | *Amborella trichopoda* | Basal angiosperms, Amborellaceae | W1P2E1 |
| Ginkgo | *Ginkgo biloba* | Gymnosperms, Ginkgoaceae | (1) |
| Cedar | *Thuja plicata* | Gymnosperms, Cupressaceae | Thupl.29378975s0017 |
| Waterfern | *Ceratopteris richardii* | Polypodiophyta (ferns), Pteridaceae | A0A8T2SGN2 |
| Common liverwort | *Marchantia polymorpha* | Liverworts, Marchantiaceae | A0A2R6XDG4 |
| | *Physcomitrium patens* | Bryophytes (mosses), Funariaceae | A0A2K1KE18 |
| Bog moss | *Sphagnum fallax* | Bryophytes (mosses), Sphagnaceae | Sphfalx03G067000 |
| Chlamydomonas | *Chlamydomonas reinhardtii* | Chlorophytes | A0A2K3DRZ9 |
| | *Auxenochlorella protothecoides* | Chlorophytes | (2) |
| | *Dunaliella salina* | Chlorophytes | Dusal.0040s00008 |

(1) https://ginkgo.zju.edu.cn/genome/blast/
(2) Fragments of exon sequence available on UniProtKB; Full length amino acid sequence from Rory J Craig, California Institute for Quantitative Biosciences (QB3), University of California, Berkeley.

**Supplementary Table 2**. **Element analysis of purified BTS-N and BTSL2-N.** ICP-MS analysis of metals co-purifying with the N-terminal domains of BTS and BTSL2 fused to MBP. The following isotopes with unique masses were quantified: $^{31}P^{16}O$, $^{54}Fe$, $^{64}Zn$, $^{65}Cu$, $^{55}Mn$, $^{60}Ni$, from which the total concentration of an element was calculated based on isotope natural abundance. Values were corrected for contaminants in the buffer and normalized to the concentration of polypeptide estimated from the sulfur ($^{32}S$) concentration.

| **Element** | **Element per protein ratio** | |
|---|---|---|
| | **MBP-BTS-N** (190 µM) | **MBP-BTSL2-N** (65 µM) |
| P | 4.16 ± 0.17 | 18.64 ± 0.54 |
| Fe | 2.69 ± 0.10 | 2.99 ± 0.05 |
| Zn | 0.31 ± 0.01 | 1.00 ± 0.01 |
| Cu | 0.06 ± 0.02 | 0.014 ± 0.00 |
| Mn | 0.29 ± 0.01 | 0.031 ± 0.00 |
| Ni | 0.04 ± 0.002 | 0.23 ± 0.00 |

**Supplementary Table 3. Primers used in the preparation of recombinant protein expression constructs for this study.**

| Protein | Forward primer | Reverse primer | Restriction enzymes used |
|---|---|---|---|
| BTSL2 N-term | ACTGCGGCCGCATGGGAGTCGGAGATCCTCTTC | TGACCTGCAGGTTATTTTTCGAACTGCGGGTGGCTCCA | NotI SbfI |
| BTS N-term | ACTGCGGCCGCATGGCGACGCCGTTACCAG | TGAGTCGACTTATTTTTCGAACTGCGGGTGGCTCCAGCATTCATTAAGCCATTCATCGAA | NotI SalI |
| BTSL2-E98A | ATAAGTATCATTCCGCAGCTGCAGATGAGGTTATATTTTCAGC | GCTGAAAATATAACCTCATCTGCAGCTGCGGAATGATACTTAT | - |
| BTSL2-E664A | CTGGAAATGCAATCTCATCTGCTGCATCTGAATGTATCTGA | TCAGATACATTCAGATGCAGCAGATGAGATTGCATTTCCAG | - |
| BTS*dgl* | CAGCACTTGGCGAAAGAAC | AAGTGCTCCGGTAGAACG | |
