## Supplementary Data 1 (SAXS data) for "Iron-sensing and redox properties of the hemerythrin-like domains of Arabidopsis BRUTUS and BRUTUS-LIKE2 proteins"

**Supplementary Data 1. SAXS experimental data and analysis.**

| (*a*) Sample details | |
| --- | --- |
| Organism | *Arabidopsis thaliana* |
| Source (Catalogue No. or reference) | Rodriguez-Celma et al., 2019 (Ref.^1^)  doi: 10.1073/pnas.1907971116 |
| Description: sequence (including Uniprot ID + uncleaved tags), bound ligands/modifications, *etc.* | Gene ID: AT1G18910  UniProt ID: F4IDY5  The protein was produced in *E. coli* with a N-terminal MBP tag which was removed by protein cleavage, leaving 9 amino acids of the linker. The protein also has a C-terminal Strep tag. The amino acid sequence is given below this table. |
| Extinction coefficient ε (wavelength and units) | 117185 M^-1^ cm^-1^ at 280 nm |
| Partial specific volume 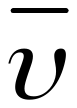 (cm^3^ g^-1^) |  |
| Mean solute and solvent scattering length densities and mean scattering contrast 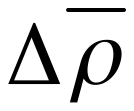 (cm^-2^) |  |
| *M* from chemical composition (Da) | 97770.91 |
| SEC SAXS Column | Superdex 200 Increase 3.2 |
| Loading volume/concentration, (mg ml^-1^) | 5.0 |
| Flow Rate (ml min^-1^) | 0.075 |
| Solvent (solvent blanks taken from flowthrough prior to elution of protein) | 10 mM MES, 15 mM NaCl, pH 6.5 |
| (*b*) SAS data collection parameters | |
| Instrument | Diamond Light Source B21 beamline with Dectris EigerX 4M detector (Ref.^2^) |
| Wavelength (Å) | 0.9464 |
| Beam geometry | 1.0 x 0.25 mm |
| Camera Length | 3.72 m |
| *q*-measurement range (Å^-1^) | 0.0045-0.34 |
| Absolute scaling method | Scaled to water scattering at 0.0163 |
| Exposure time (s) | 3.0 |
| Capillary Diameter (mm) | 1.5 |
| Sample temperature (ºC) | 15 |
| (*c*) Software employed for SAS data reduction, analysis and interpretation | |
| SAS data reduction to sample–solvent scattering, and extrapolation, merging, desmearing *etc*. as relevant | Solvent subtraction and merging performed using ScÅtter [https://sibyls.als.lbl.gov/scatter/] version IV.d |
| Calculation of ε from sequence |  |
| Calculation of 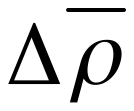 and 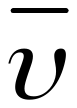 values from chemical composition |  |
| Basic analyses: Guinier, *P*(*r*), scattering particle volume (*e.g.* Porod volume *V*_P_ or volume of correlation *V*_c_) | PRIMUS from ATSAS 3.2.1 (Ref.^3^) |
| Shape/bead modelling | DAMAVER Suite (Ref.^4^) via PRIMUS from ATSAS 3.2.1; GASBOR via ATSAS online (https://www.embl-hamburg.de/biosaxs/atsas-online) |
| Atomic structure modelling (homology, rigid body, ensemble) | Alphafold (Ref.^5^), SREFLEX (Ref.^6^) |
| Molecular graphics | Chimera X (Ref.^7^) |
| (*d*) Structural parameters | |
| Guinier Analysis | |
| *I*(0) (cm^-1^) | 0.062 ± 0.000096 |
| *R*_g_ (Å) | 33.95 ± 0.09 |
| *q-*range (Å^-1^) | 0.005-0.235 |
| sRg Limits | 0.28, 1.30 |
| *M* from *I*(0) (ratio to expected value) | 1.03 |
| *P*(*r*) analysis | |
| *I*(0) (cm^-1^) | 0.06 |
| *R*_g_ (Å) | 33.58 |
| *d*_max_ (Å) | 96.79 |
| *q-*range (Å^-1^) | 0.005-0.235 |
| Total quality estimate | 0.83 |
| *M* from Bayesian Inference (ratio to expected value) | 101050 (1.03) |
| (*e*) Shape modelling results | |
| DAMMIN | |
| *q-*range for fitting (Å^-1^) | 0.005-0.235 |
| Symmetry/anisotropy assumptions | P1 |
| Ambiguity measure(s) with definitions |  |
| χ^2^ value | 1.075 |
| *P* value, any other quality-of-fit parameters | 0.051448 |
| Resolution from SASRES (Å) | 1000 |
| Relevant output parameters (*e.g.* predicted *R*_g_/*d*_max_ values, weights for multi-state models, *etc.*) |  |
| Domain/subunit coordinates and contacts, regions of presumed flexibility as appropriate |  |
| (*f*) Atomistic modelling | |
| Alphafold Model | Rank 2 |
| *q-*range for modelling (Å^-1^) |  |
| CRYSOL |  |
| χ^2^, *P*-Value | 1.008, 0.04 |
| (*g*) SASBDB IDs for data and models | |
| Data | www.sasbdb.org/data/SASDU79/1g65egyk1i/ |

**Amino acid sequence of cleaved BTSL2-NT,** calculated mol. weight: 97770.91 Da

ISHMSMGGRMGVGDPLPLPPEKNRREVNKPPDIASTSSSSASAVNNARLSDAPILLFVYFHKAFRAQLAELQFLAGDTVRSGSDLAVELRSKFEFLKLVYKYHSAAEDEVIFSALDTRVKNIVFNYSLEHDATDDLFTSVFHWLNVLEEEQGNRADVLREVVLCIGTIQSSICQHMLKEERQVFPLMIENFSFEEQASLVWQFICSVPVMVLEEIFPWMTSLLSPKEKSEVETCFKEVVPNELSLQLVINSWLIDDSQSSLTALTKIMKGVQSVEVSENMTNSQTNSSSSGVFQRFWQWSKKMSFSSPNTGHILVHGIHLWHNAIRKDLVDIQKGLCQLTFPSLSLDLNVLVVRLNFLADVLIFYSNAFKTFFYPVFEDMVDQQHSSSSKQFTIDGHVENFKKSLDLETRAGSDNFVITLQEKLESLILTVAKQFSIEETEVFPIISKNCNIEMQRQLLYRSIHFLPLGLLKCVIMWFSAQLPEDECQSIIHYLSSEDSFPNKPFAHLLLQWFRFGYSGKTPVESFWNELSFMFKPRCSFEEELTEEASGSFFQQSPQKLFKVSDPYSMDPPAGYMNETPYSSAMNQQILIPGKLRPLLHLPDLFGDKTIGEHLTMDLKPIDLIFYFHKAMKKDLDYLVRGSARLATDYSFLGEFQQRFHLIKFLYQIHSDAEDEIAFPALEAKGKLQNISQSYSIDHELEVEHLNKVSFLLNELAELNMLVLDHKNVKYEKLCMSLQDICKSIHKLLSEHLHREETELWCLFRDCFTIEEQEKIIACMLGRISGEILQDMIPWLMESLIPDEQHAVMSLWRQATRKTMFGEWLTEWYNSHAVEEETEEAWSHPQFEK
